# Pupillary dynamics reflect age-related changes in memory encoding

**DOI:** 10.1101/2025.10.23.683937

**Authors:** Adrian Ruiz Chiapello, Enzo Buscato, Muriel Mescam, Isabelle Berry, Alexandra Pressigout, Andrea Alamia, Florence Remy

## Abstract

Pupillary dynamics are closely dependent on tonic and phasic activity of the brainstem locus coeruleus (LC), a key neuromodulatory nucleus. LC neurons are the earliest site of hyper-phosphorylated tau accumulation, leading to a decline in nucleus structural integrity in older age that likely impacts long-term memory. While several studies have explored the link between pupil dilation and successful memory encoding, little is known about the effects of aging on this relationship. This study investigated task-evoked pupillary responses (TEPRs) in young and older adults during incidental encoding of neutral visual scenes. Memory performance was assessed 24 hours later using a Remember-Know-Guess recognition task. TEPRs were compared based on recognition performance. Our findings notably revealed attenuated TEPRs in older individuals, supporting the hypothesis of impaired LC-driven modulation with age. Pupil dilation was associated with memory performance in young adults only. In this group, subsequently recognized stimuli elicited greater dilation during encoding compared to forgotten ones. Moreover, among recognized stimuli, recollected ones evoked larger pupillary responses than those remembered with a feeling of familiarity. Importantly, this effect was absent in older adults, suggesting that the benefit provided by LC involvement during encoding declines with advancing age. These findings highlight the crucial role of LC-mediated neuromodulation in episodic memory, and suggest that age-related LC decline can be evidenced using pupillometry.

- Evoked pupillary dilation and tonic pupil size were both decreased in aging.
- Increased pre-stimulus pupil diameter was associated with altered stimulus processing.
- Pupillary dilation at encoding was greater for later recognized items in young adults.
- Pupillary dilation at encoding was not related to subsequent memory in older adults.
- In aging, reduced LC involvement at encoding may lead to lower memory performance.

## 1. Introduction

Aging is associated with a progressive decline in specific cognitive abilities, including executive functions (Salthouse, 2010) and episodic memory (Nyberg et al., 2012; Rönnlund et al., 2005). Activity in brain networks underlying these cognitive processes is largely influenced by neuromodulatory inputs (Shine et al., 2019). In particular, ascending projections from the locus coeruleus (LC) (Szabadi, 2013) release noradrenaline (NA) to cerebral regions, allowing optimization of executive functions (Poe et al., 2020; Chandler et al., 2014), and encoding, consolidation, and retrieval phases of long-term memory (Dahl et al., 2023; Sara, 2015, Sara, 2009). The LC-NA system evidences early alterations with age (Braak and Del Tredici, 2011), which may thus partly account for the selective impairment in executive and memory performance in older individuals (Mather and Harley, 2016). Since LC activity has been tightly linked to pupil dynamics (Joshi and Gold, 2020), pupillometry is considered a relevant tool for investigating how noradrenergic modulation influences cognitive performance, and how this influence evolves in aging (Elman et al., 2017; Mather and Harley, 2016). In the present study, we aimed to explore the relationship between pupil dynamics and performance in healthy young and older adults, using implicit encoding of visual scenes and their long-term recognition.

The LC exhibits two modes of neural activity, tonic and phasic, to modulate cerebral excitation or inhibition. According to the adaptive gain theoretical framework (Aston-Jones and Cohen, 2005), transient bursts of phasic LC activity optimize task-related performance, as temporary NA release in brain regions involved in the task enhances selectivity of activations (Mather et al., 2016) for optimal performance. In addition, tonic LC activity levels are thought to influence performance. While optimal baseline activity promotes task engagement, lower levels are linked to reduced alertness. Conversely, higher tonic activity facilitates external exploration in changing environments to support behavioral updating, though it may also heighten distractibility (McGinley et al., 2015; Sara and Bouret, 2012). Throughout adulthood, LC neurons are the very first to show intra-cellular accumulation of hyper-phosphorylated tau proteins (Harley et al., 2021; Braak and Del Tredici, 2011). These tau aggregates are thought to occur decades before any neuronal loss, and to result in altered tonic and phasic LC activity, impacting NA-dependent cognitive abilities (Weinshenker, 2018; Mather and Harley, 2016). A recent study in rodents (Kelberman et al., 2023) reported transient LC phasic hyperactivity with abnormal tau accumulation in the nucleus, followed by LC hypoactivity as tau pathology spreads to brain regions. In the latter rodent model, LC tau aggregates were also associated with a decrease in LC tonic activity, although an age-related increase of LC tonic discharges has been suggested in humans (Ehrenberg et al., 2018; Weinshenker, 2018; Elman et al., 2017). In vivo assessment of LC integrity is possible using magnetic resonance imaging (MRI), through measures of LC neuromelanin content (Betts et al., 2017). Intracellular neuromelanin is a by-product of NA metabolism, and neuromelanin content is dependent on NA levels and on the number of LC neurons (Beardmore et al., 2024). Human neuroimaging studies have consistently shown a decrease in LC integrity from the age of 60 in healthy participants (Liu et al., 2019; Betts et al., 2017; Shibata et al., 2006), and a link between LC integrity and cortical tau pathology (Jacobs et al., 2021). MRI-assessed LC integrity has also been related to standard neuropsychological scores (Jacobs et al., 2021; Liu et al., 2020) and to working memory (Elman et al., 2017) and emotional episodic memory performance (Hammerer et al., 2018). Altogether, this converging evidence underlines early LC-NA system dysfunction with age, its impact on cognition, and the relevance of in vivo LC biomarkers.

Several studies have consistently shown that pupil dynamics are closely correlated with LC neuronal activity, recorded either using electrophysiology in rodents (Breton-Provencher and Sur, 2019; Liu et al., 2017; Reimer et al., 2016; Carter et al., 2010) and non-human primates (Joshi et al., 2016; Varazzani et al., 2015), or using functional MRI in humans (De Gee et al., 2017; Murphy et al., 2014). In young individuals, tonic and phasic pupil changes have been respectively correlated with resting-state (Wu et al., 2025; DiNuzzo et al., 2019) and task-evoked (Clewett et al., 2018; Murphy et al., 2014) fMRI LC activity. Two major noradrenergic pathways are implicated in the control of pupil diameter by the LC. Firstly, LC projections to the spinal cord stimulate the orthosympathetic pathway that controls pupil dilation. Secondly, LC projections to the Edinger-Westphal nucleus (EWN) inhibit the parasympathetic pupil constriction reflex loop (Joshi and Gold, 2020; Larsen and Waters, 2018; Mathôt, 2018). In relation to LC activation dynamics, tonic pupil size is thought to reflect basal vigilance states and sustained attentional control, while phasic dilation amplitude may result from transient arousal due to increased alertness, emotion, or cognitive engagement in response to stimuli. Accordingly, relationships between pupil and cognitive measures have been extensively reported in young adults. Pupil dilation has been linked to executive functions (van der Wel and van Steenbergen, 2018; Unsworth and Robison, 2017), such as inhibitory control (Rondeel et al., 2015; Wang et al., 2015; Laeng et al., 2011), cognitive flexibility (Rondeel et al., 2015), and working memory (Brouwer et al., 2014; Piquado et al., 2010). Pupillometry studies using memory encoding paradigms have provided mixed results. While some studies have linked successful encoding in long-term memory with increased phasic pupil dilation (Pitem and Mama, 2025; Bergt et al., 2018; Kucewicz et al., 2018; Papesh et al., 2012), others have reported an association with either pupil constriction (Pilarczyk et al., 2022; Naber et al., 2013), reduced pupil dilation (Kafkas and Montaldi, 2011) or no association with pupil dynamics (Cronin et al., 2023; Hammerer et al., 2017; Võ et al., 2008). Pre-stimulus tonic pupil diameter may also condition subsequent encoding performance (Pitem and Mama, 2025; Cronin et al., 2023; Kucewicz et al., 2018; Hammerer et al., 2017). Here, the discrepancy between studies may likely rely on differences in experimental paradigms, and specifically in encoding instructions.

In line with age-related alterations in noradrenergic neuromodulation, changes in pupil dynamics have been observed in older individuals. Average tonic pupil size decreases linearly with age (Huang et al., 2024; Tekin et al., 2018), and pre-stimulus tonic diameter may remain a determinant of visual task performance in older subjects (Moran et al., 2025). Studies of phasic pupil response during task completion in older subjects are relatively sparse, and the cognitive correlates of age-related pupil changes remain unclear. Phasic pupil dilation was observed in older adulthood for various cognitive tasks, and its amplitude was generally reported to decline with age (Riley et al., 2024; He et al., 2020; Lee et al., 2018; Hammerer et al., 2017; Van Gerven et al., 2004), thus indicating weakened phasic LC activity. It should be noted that no agerelated changes in pupil size were observed during simple motor reaction time tasks (Ribeiro and Castelo-Branco, 2019), and that age-related reductions in pupil responses were more evident under higher task difficulty (Granholm et al., 2017; Van Gerven et al., 2004). Regarding links between pupil dynamics and encoding in long-term memory, Hammerer et al. (2017) reported an age-related reduction in phasic dilation during encoding of negative compared to neutral visual scenes, with no relationships between pupil dilation and subsequent memory performance in any age groups. In agreement, Lee et al. (2018) found that an emotionally-arousing context of encoding elicited smaller pupil dilation in older versus young subjects. However, subsequent stimulus memory was not tested in the latter study. To our knowledge, there is no report yet on age effects on pupil responses during non-emotional encoding in long-term memory, and on the relationship with subsequent recognition and recollection, known to be both impaired in aging (Koen and Yonelinas, 2014). Since the latter relationship has been reported in young adults, although with mixed results (Papesh et al., 2012; Kafkas and Montaldi, 2011), we aimed at exploring it in healthy older adults, to investigate possible LC-driven benefit for successful memory encoding.

In the present study, pupil dynamics were recorded in young and older individuals during a fixation task, allowing measurement of basal pupil dynamics at rest, and a rapid visual object categorization task. The latter task featured neutral scenes containing salient animal and furniture objects within congruent or incongruent backgrounds (Rémy et al., 2013). Due to brief scene presentation, this task required high sustained engagement and selective attention to objects in scenes, both processes being dependent on LC tonic and phasic activity (McGinley et al., 2015; Sara, 2009). Moreover, incongruent scenes were expected to capture higher attention and trigger greater cognitive effort for conflict resolution, therefore possibly enhancing LC-driven arousal and pupil response (Grueschow et al., 2021; Köhler et al., 2016). The categorization task served as an incidental encoding phase for a recognition memory test conducted 24 hours later. We investigated whether pupil dynamics at encoding could predict immediate success in the categorization task, as well as subsequent long-term scene recognition and memory strength. Pupil results were interpreted in terms of LC involvement in these tasks, along with age-related effects that may indicate LC-NA system dysfunction. Building on the studies mentioned earlier, we anticipated an age-related effect on both baseline and task-evoked pupil dynamics, with older participants exhibiting reduced pupil size and phasic dilation. In young adults, we expected variability in phasic pupil responses to various visual scenes, with greater pupil dilation promoting successful stimulus encoding. Additionally, we anticipated that subsequent memory strength would depend on stimulus-evoked pupillary response during encoding. In older adults, alterations in LC tonic and phasic activity may reveal diminished involvement of the LC-NA system in supporting effective stimulus encoding. Accordingly, pupil dynamics at encoding may no longer predict subsequent long-term recognition in older age.

## 2. Materials & Methods

### 2.1. Participants

A total of 112 participants were recruited for this study. Data from 4 participants (3 younger, 1 older) were removed due to defective measurements. Consequently, the final sample comprised 108 participants, of which 57 were younger individuals (34 female, aged between 18 and 30, mean age ± SD = 22.2 ± 2.2 years) and 51 were older individuals (35 female, aged between 58 and 88, mean age ± SD = 70.8 ± 6.9 years). Gender distribution was equivalent between groups (*p* = 0.332). All participants had normal or corrected-to-normal vision: they scored at least 8.2 out of 10 on the Armaignac test (visual acuity) and 1.50 out of 1.95 on the Pelli-Robson test (contrast vision). None of them had a history of any neurological or psychiatric disorders, and all had a Mini-Mental State Examination (MMSE) greater than or equal to 27. Number of education years was slightly lower in the older group (mean ± SD = 15.7 ± 1.4 in the younger group; mean ± SD = 14.4 ± 3.6 in the older group; *p* < 0.01). Informed written consent was obtained for every participant. The study was approved by the French ethics committee CPP Sud-Ouest et Outre-Mer I (ID-RCB 2021-A02686-35).

### 2.2. Stimuli

The stimuli were taken from a large set of visual scenes described in a previous report (Rémy et al., 2013). We used a set of 144 real-life color pictures, in which a foreground object, either an animal or a piece of furniture (n_*animal*_ = n_*furniture*_ = 72), was superimposed on either an indoor or outdoor context (n_*indoor*_ = n_*outdoor*_ = 72). Congruent stimuli were defined as scenes featuring an animal in a natural outdoor environment, or a piece of furniture in an artificial indoor environment, e.g., a polar bear on an Arctic ice pack or a chair in a room. On the other hand, incongruent stimuli were defined as scenes featuring an animal in an artificial indoor environment or a piece of furniture in a natural outdoor environment. There were as many congruent stimuli as incongruent ones (n_*congruent*_ = n_*incongruent*_ = 72). All stimuli were equalized in contrast and luminance. The average object size was 12.7 ± 4.7% of the whole image. Examples of stimuli are shown in Fig. 1. The set of stimuli was divided into 2 subsets of 72 scenes, with an equal number of congruent and incongruent scenes in each subset. Numbers of animals and pieces of furniture were also equalized in each subset, as well as numbers of outdoor and indoor contexts.

**Figure 1.**
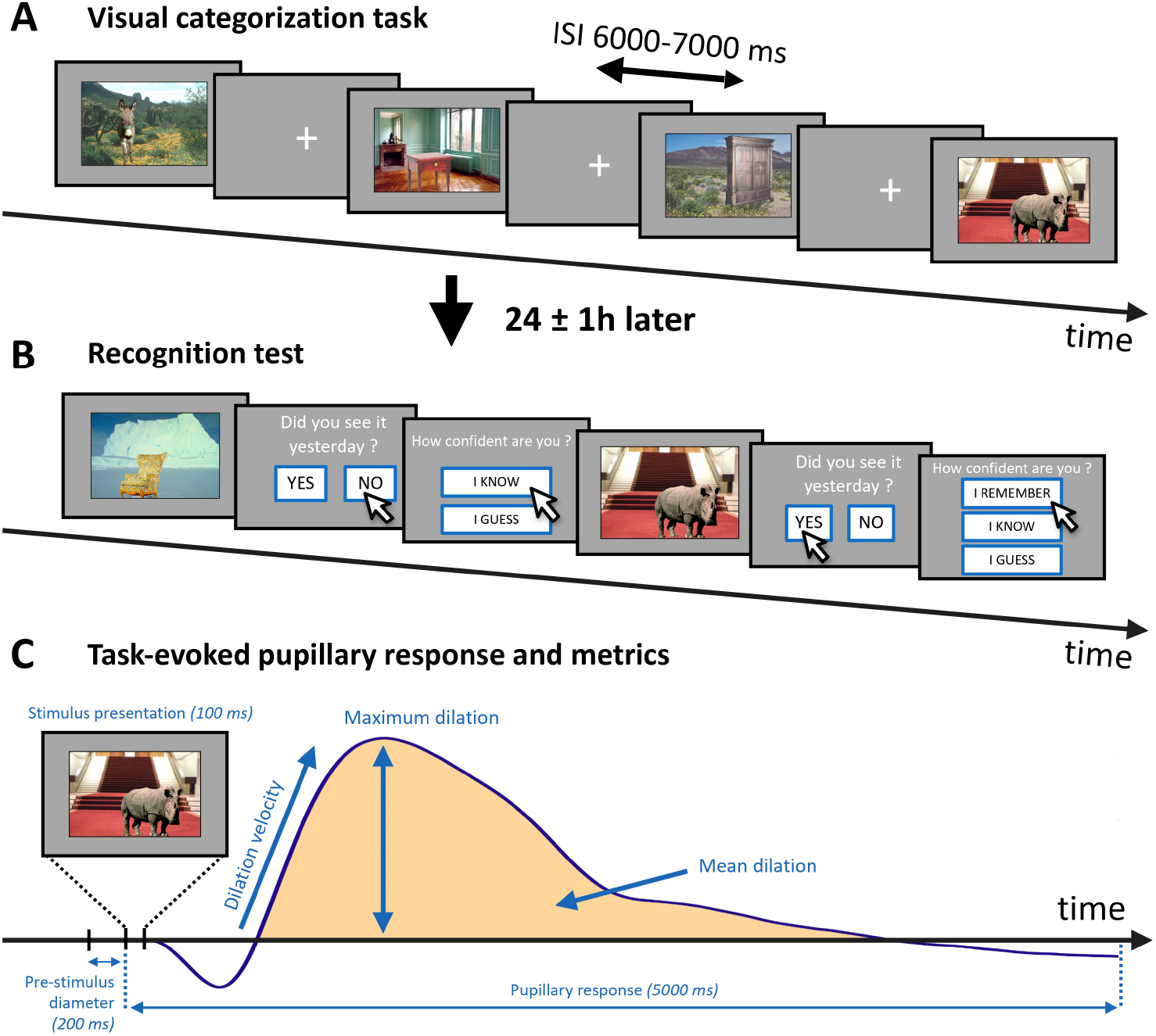
Experimental procedure. (**A**) Encoding session. Each scene was presented during 100 ms, with an inter-stimulus interval (ISI) ranging from 6000 ms to 7000 ms. Foreground objects in scenes had to be categorized as accurately and as fast as possible (animal or piece of furniture). (**B**) Recognition test. Scenes were presented during 4 s. Participants had to recognize previously encountered scenes among novel scenes, and rated their confidence (Remember, Know, or Guess). (**C**) Task-evoked pupillary response and extracted metrics. The response averaged across all participants and all trials is shown here for illustrative purposes.

### 2.3. Experimental procedure

Volunteers were asked to take part in two experimental sessions on two consecutive days. Across all participants, the average interval between both sessions was 24 ± 1h (mean ± SD). Pupil diameter was recorded only during the first session.

#### 2.3.1. First experimental session

The first session consisted of a resting task, followed by a fast visual categorization task. Participants were seated on a chair in a quiet and dimly-lit room with the head stabilized by a chin rest, in front of a 22-inch monitor placed 60 cm away from their eyes. Pupil dynamics were recorded with a sampling frequency of 300 Hz, using the Eyebrain T2 Tracker (SuriCog, Paris, France). Volunteers were asked to blink as little as possible during the tasks. The eye-tracker was calibrated before pupil measurements, using a standard 13-point calibration.

The session started with the resting task, during which participants had to fixate a central cross on the screen for 2 minutes. Then, the fast visual categorization task was conducted, using a subset of 72 visual environmental scenes, as described above. Each trial started with a fixation cross that was displayed during a random interval between 6000 and 7000 ms, followed by a scene briefly presented during 100 ms (Fig. 1A). Stimulus presentation was made short in order to prevent eye movements and subsequent pupil foreshortening errors. Participants had to categorize the foreground object in the scene as either an animal or a piece of furniture, as quickly and accurately as possible. To respond, they had to press one of two available keys on a keyboard with their dominant hand. The key side and stimulus subsets were randomly assigned to participants and counterbalanced. The 72 trials were divided into 4 blocks of 18 trials each. Each block lasted approximately 2 minutes. A break of variable duration (managed by the participant) was made between each block. Importantly, participants were not informed that they would have a recognition test the following day. Therefore, the fast categorization task enabled incidental encoding of the 72 presented scenes.

### 2.3.2. Second experimental session

The second session consisted of a surprise scene recognition task. Participants were presented with 144 stimuli, including the 72 previously encoded scenes intermixed with the 72 novel scenes of the other subset. In a trial, a unique scene was displayed for 4000 ms, then participants had to indicate whether the scene had been presented or not during the categorization task. They answered two questions (Fig. 1B): first, “Did you see this image yesterday?” with a *yes* or *no* response, and second, a confidence judgment assessing their subjective experience of recognition. This was based on the Remember-Know-Guess (RKG) paradigm (Tulving, 1985), which required participants to indicate whether their recognition was based on detailed recollection of contextual elements (R for *remember*), a feeling of familiarity (K for *know*), or a mere guess (G for *guess*).

### 2.4. Data analyses

Automatic pipelines made in-house using MatLab 2021a were used for pupil raw signal pre-processing, pupil time-series comparisons between groups and conditions, and computation of pupil response metrics (code and graphical interface available at https://github.com/Adrian-R-C/pupil-study-1-code). Statistical analyses of behavioral and pupil-derived variables were conducted using R v.4.4.3 and Jamovi v.2.3.28.

#### 2.4.1. Behavioral data

Global recognition performance for each participant was defined as the number of correct answers, i.e., the sum of subsequently recognized targets (*N*_*hits*_) and correctly rejected distractors (*N*_*correctly rejected*_), divided by the total number of test stimuli (*N*_*test*_ = 144). Participants’ recognition was above chance level if performance was greater than or equal to 59 % (binomial distribution for 144 trials, *α* = 0.05). Individual indices of recollection (R), familiarity-based recognition (K), and guessing (G) were calculated by subtracting the *false alarm rate* from the *hit rate* for each confidence level.

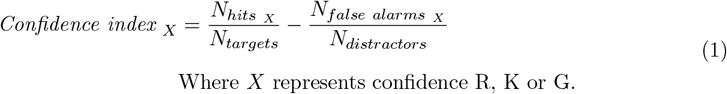

with 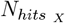 and 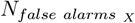 being respectively the number of recognized targets and false alarms with confidence *X, N*_*targets*_ = 72 the number of previously encountered targets, and *N*_*distractors*_ = 72 the number of novel stimuli. These three confidence indices together help discriminate the contributions of different retrieval processes to overall recognition performance (derived from Yonelinas (2002); Tulving (1985)).

Global categorization and recognition performance were compared across age groups by computing independent t-tests. Additionally, the effects of stimulus properties on categorization (in terms of accuracy and response time) and recognition performance were examined using two-way ANOVAs, incorporating age group as a between-subjects factor and either scene congruence (congruent vs. incongruent) or object type (animal vs. furniture) as a within-subjects factor. Finally, the influence of age on confidence indices (Eq. 1) in the recognition task was explored using a two-way ANOVA, with factors of age group (inter-subject) and confidence level (intra-subject, R vs. K vs. G). Two-tailed post-hoc tests were conducted, and p-values were corrected using the False Discovery Rate (FDR) correction method.

#### 2.4.2. Pupil data pre-processing

For both fixation and categorization tasks, horizontal and vertical pupil diameters were continuously measured from both eyes. Only horizontal diameters were used for analysis, since they were less affected by blinks, and were averaged between left and right eyes. Blinks were identified by thresholding the 1^*st*^-order time derivative of pupil diameter. The threshold was set empirically. Portions of the signal affected by blinking artifacts, as well as 50 ms periods before and after the blink, were removed and linearly interpolated. Hampel detection was applied to locate outliers, with a number of adjacent samples of *K* = 5 and a normalized deviation threshold of *N*_*sigma*_ = 2, and time outliers were replaced with the local median (Leys et al., 2013). The signal was finally detrended and low-pass filtered at 4 Hz (Kret and Sjak-Shie, 2019).

#### 2.4.3. Basal pupil measures

The mean basal pupil diameter was computed from the 2-minute resting task by taking the average value of the pupil signal, skipping the first second due to monitor luminance variation. Basal diameters were compared across age groups with a one-tailed independent t-test, under the hypothesis that younger individuals would exhibit a larger basal pupil diameter. Spontaneous fluctuations amplitude was estimated using the following formula: (SD/Mean) · 100, expressed in % of the mean basal pupil diameter (Aminihajibashi et al., 2019). Fluctuation amplitude was compared between younger and older participants using a two-tailed Welch t-test to account for unequal variance.

#### 2.4.4. Task-evoked pupillary response

Pupillary signal from the categorization task was split into 72 segments, with each segment onset corresponding to trial onset (stimulus at *t* = 0*s*; Fig. 1C). Pupil data from each trial were baseline-corrected by subtracting the mean signal over the 200-ms pre-stimulus period. Trials with excessively interpolated data, i.e., more than 25% of trial duration, were excluded.

The median pupil time series for each participant were computed on a 5-second epoch starting from stimulus onset. Median task-evoked pupil responses (TEPRs) over all trials were compared between age groups to assess age effects on the phasic pupil response. Within-subjects differences between task conditions were evaluated in each age group and for each experimental factor (categorization success, recognition success, recognition confidence level, scene congruence, and object type). For all comparisons, subjects’ median TEPRs were not included if there was only 6 valid trials or less to compute the median (trials with interpolation below 25% were considered valid). In all within-subjects analyses conducted separately in each age group, TEPRs were expressed as an absolute variation in diameter following a stimulus, since pre-stimulus diameter and phasic dilation amplitude were not correlated (see Results). Regarding comparisons between age groups, both absolute (signal − baseline) and relative 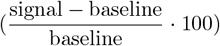 variations in diameter were considered.

To test for significant differences between conditions in the pupil time series, we implemented a threshold-free cluster enhancement (TFCE) procedure (Mensen and Khatami, 2013; Smith and Nichols, 2009). This approach does not rely on a predefined cluster threshold, unlike conventional cluster-based permutation testing. At each time point, either paired or independent t-tests were computed. Raw t-values were then transformed into TFCE scores, which combine both the test statistic magnitude (height, weighted by parameter *H* = 2) and the temporal support from contiguous data points (extent, weighted by parameter *E* = 1). Significance was assessed via permutation testing: condition or group labels were randomly shuffled 5,000 times, and for each permutation, new TFCE scores were generated to form a null distribution of TFCE scores. Computed TFCE scores in unpermuted data were then compared against this null distribution to obtain corrected p-values, ensuring control of the family-wise error rate across time points. The resulting p-values were thresholded at *α* = 0.05. We report the mean t-statistic and mean Cohen’s d as overall estimates of effect size, averaged across all time points that reached significance. Uncorrected results remain accessible for exploration through our GUI (https://github.com/Adrian-R-C/pupil-study-1-code).

In addition to the temporal analysis of TEPRs, four pupillary metrics were derived from the pupillary signals (Fig. 1C): *pre-stimulus pupil diameter*, calculated as the mean pupil size during the 200 ms preceding stimulus onset; *dilation amplitude*, defined as the maximum pupil diameter within the 5-second response window; *dilation velocity*, computed as the dilation amplitude divided by the time to peak; and *mean dilation*, representing the average pupil dilation over the 5-second response period. The variability of pupil phasic responses in each participant was calculated using the standard deviation of the dilation amplitude across all 72 trials, relative to the average dilation amplitude. This variability was expressed using the formula: (SD/Mean) · 100. Pupillary metrics were further analyzed on a subject and trial basis, as follows.

In a subject-based approach, median phasic features in each participant were compared between conditions and age groups using two-way ANOVAs. Two-tailed or one-tailed (when specified, based on a priori assumptions) post-hoc tests were conducted, and p-values were corrected using FDR correction. The link between median pupil dilation at encoding and global recognition performance was assessed with Spearman correlation coefficient. Moreover, we explored the links between median phasic dilation amplitude and median pre-stimulus diameter, and between mean resting diameter and median pre-stimulus diameter. Correlations were investigated across all individuals and separately in each age group, and p-values were corrected using FDR correction.

In a trial-based approach, we used generalized linear mixed models (GLMMs) to explore the relationships between pupillary metrics and cognitive performance, and the influence of stimulus properties on the pupillary response. In each of the models, a random intercept was included to account for individual variability in cognitive performance.

A first set of models explored the relationship between pupillary metrics and categorization success. The influence of pre-stimulus pupil diameter was assessed using a GLMM with trial’s categorization success as the binary dependent variable, and age, pre-stimulus pupil diameter, and their interaction as explanatory regressors.

A model, including additional fixed factors, i.e., scene congruence and object type, was also implemented. Two similar models were constructed to examine the relationship between stimulus-evoked dilation and categorization success, with the latter as the binary dependent variable, and age, dilation amplitude, and their interaction as explanatory regressors.

A second set of models was implemented to explore the relationship between pupillary measures and recognition success, in which categorization success was replaced by subsequent recognition success as the binary dependent variable. These models included only participants who performed above chance level on the recognition task (see memory performance results).

Finally, the effects of stimulus properties on pupillary dilation were also investigated in a trialwise model. Here, dilation amplitude was the dependent variable, and congruence, foreground object type, and their interaction were the fixed factors.

## 3. Results

### 3.1. Global age effect on resting pupil and evoked pupillary response

In total, 101 subjects from the resting task could be included in the analysis (53 younger and 48 older participants). Mean pupil basal size was significantly lower in older compared to younger participants (*t*(99) = 3.846, *p* < 0.001, Cohen’s *d* = 0.766, one-tailed; Fig. 2A, left panel). Pupil fluctuations amplitude was also lower in older relative to younger participants (*t*(87.34) = 7.280, *p* < 10^−9^, Cohen’s *d* = 1.418; Fig. 2A, right panel).

**Figure 2.**
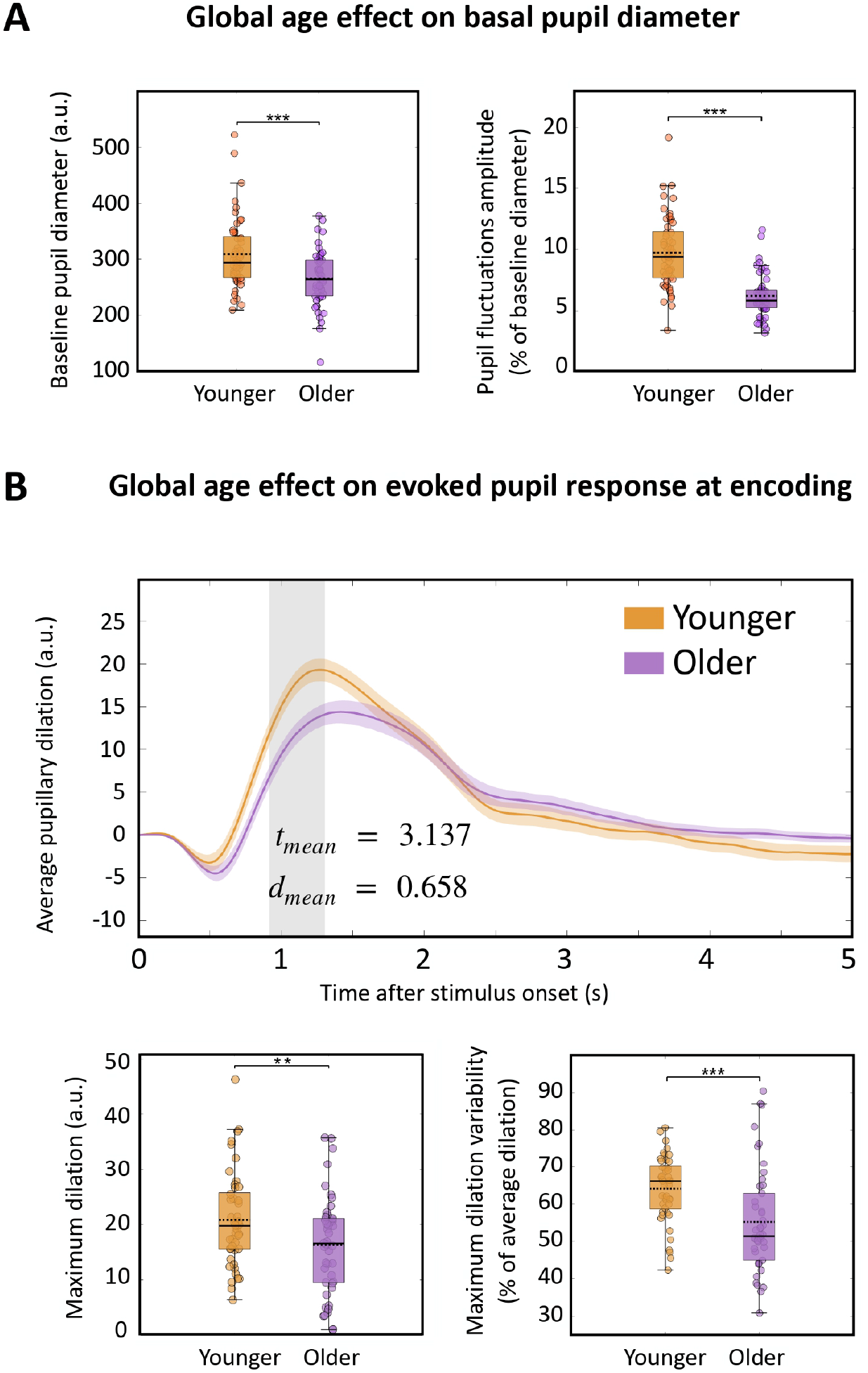
Age effects on basal and evoked pupil diameter. (**A**) Fixation task. Resting-state average pupil diameter and pupil fluctuations in younger and older individuals. (**B**) Categorization task. Upper panel: TEPRs averaged across participants in younger (*n* = 45) and older (*n* = 46) groups. Shaded gray area represents significant difference between groups (one-tailed independent t-tests with *H*_1_ : younger *>* older, TFCE-corrected cluster with *p* < 0.05). Error bar represents s.e.m. Lower panels: average peak dilation amplitude and peak dilation variability across trials in both age groups. (^∗^*p* < 0.05;^∗∗^ *p* < 0.01;^∗∗∗^ *p* < 0.001, FDR correction). The solid and dotted lines in boxplots represent group median and mean values, respectively.

For the categorization task, 91 pupil signals could be included in the analysis (45 younger and 46 older participants). Eleven participants were omitted due to technical issues, and pupil data from 6 individuals were removed as a large proportion of their trials contained excessive missing data. Increased evoked pupil dilation was observed in the younger relative to the older group (mean t = 3.137, mean Cohen’s *d* = 0.658, TFCE-corrected; Fig. 2B, upper panel). When considering the response as a percentage of dilation relative to the pre-stimulus diameter, a similar cluster was initially observed but did not survive TFCE correction (figure not shown here). Maximum dilation and dilation velocity were greater in younger participants (maximum dilation: *t*(89) = 2.454, *p* = 0.009, Cohen’s *d* = 0.515, one-tailed; Fig. 2B, bottom left; velocity: *t*(89) = 3.128, *p* = 0.002, Cohen’s *d* = 0.656, one-tailed). The difference in mean dilation between groups was not statistically significant (*p* = 0.131). Our results also revealed that younger individuals showed higher trial-to-trial dilation variability (mean ± SD = 64.2 ± 9.0 % of average dilation) than older adults (55.3 ± 14.0 %; *t*(76.9) = 3.622, *p* < 0.001, Cohen’s *d* = 0.756, Fig. 2B, bottom right).

Median pre-stimulus pupil diameter measured during the categorization task correlated with mean pupil diameter at rest over all participants (*ρ*(84) = 0.785, *p* < 0.001; Fig. S1A), and also in both group separately (younger: *ρ*(41) = 0.721, *p* < 0.001; older: *ρ*(41) = 0.824, *p* < 0.001). Moreover, pre-stimulus pupil diameter was not correlated with phasic dilation amplitude, when considering all participants (*ρ*(89) = 0.170, *p* = 0.108; Fig. S1B) or within each age group (younger: *ρ*(43) = 0.219, *p* = 0.149; older: *ρ*(44) = 0.045, *p* = 0.764). This shows importantly that individuals’ amplitude of phasic response to the stimulus did not depend on pre-stimulus basal diameter.

### 3.2. Pupil dynamics and categorization performance

Global accuracy on the categorization task was high in both groups, although reduced in older (mean ± SD = 82.1 ± 13.2 %) relative to younger adults (95.4 ± 5.4 %; *t*(60.4) = 6.273, *p* < 10^−6^, Cohen’s *d* = 1.302; Fig. 3A, left panel). Moreover, mean response times for correct trials (RTs) were longer in the older group (mean ± SD = 701 ± 123 ms) compared to younger counterparts (637 ± 84 %; *t*(79.6) = 2.896, *p* = 0.006, Cohen’s *d* = 0.606; Fig. 3A, right panel). Importantly, categorization errors were attributable to time pressure rather than an inability to perceive the scenes, as participants were most often aware and reported their own categorization mistakes.

**Figure 3.**
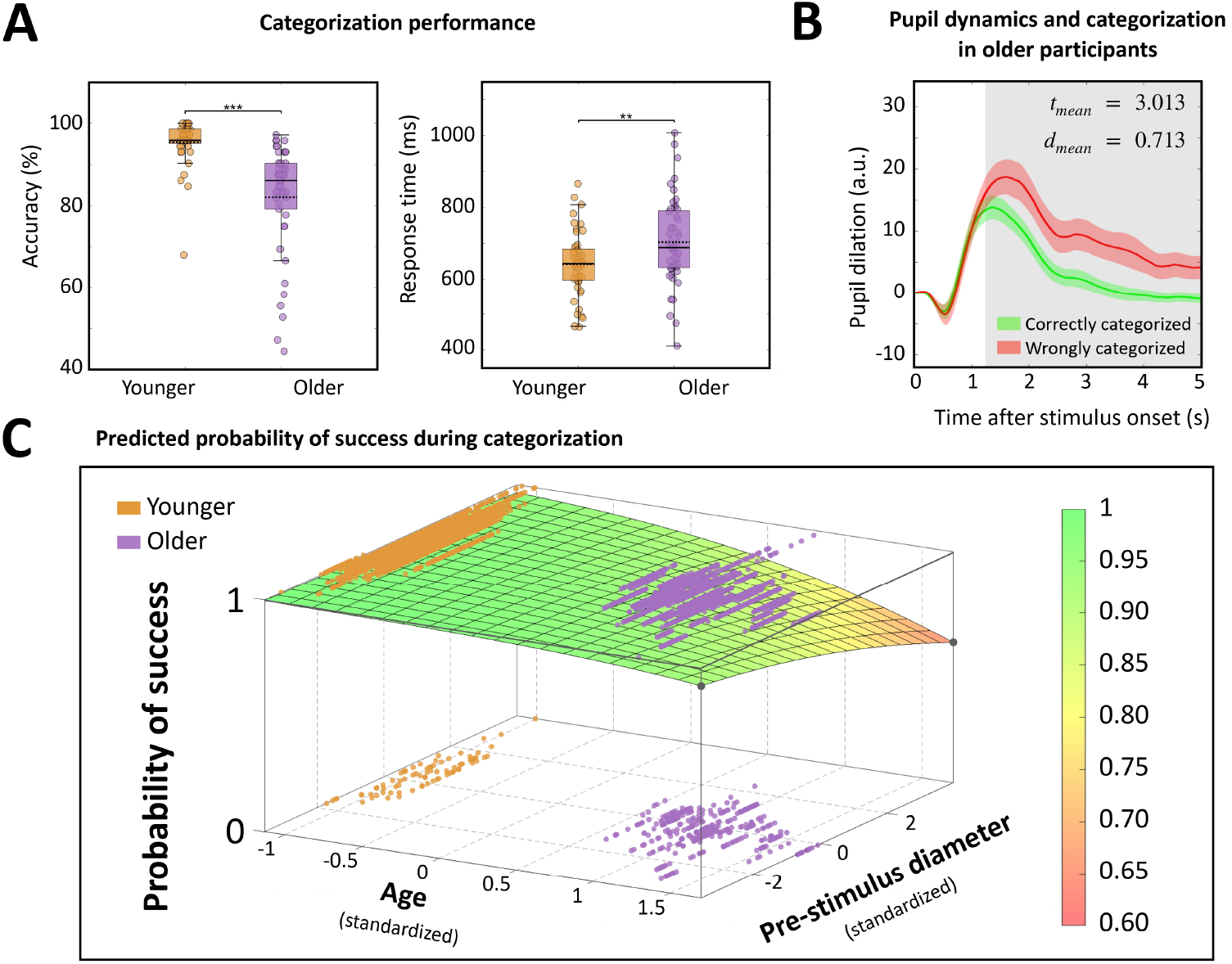
Categorization results. (**A**) Accuracy and mean response time on correct trials in both age groups. (**B**) TEPRs in older individuals (*n* = 20) for correct and incorrect responses. Shaded gray area represents significant difference between conditions (two-tailed paired t-tests with *H*_1_ : correctly categorized *≠* wrongly categorized, TFCE-corrected cluster with *p* < 0.05). Error bar represents s.e.m. (**C**) Predicted probability of success as a function of age and pre-stimulus pupil diameter. The model predicts the probability according to the following formula: *P* (success) = (1 + *exp*(−*β*_0_ − *β*_1_ · *Age* − *β*_2_ · *Pre-stimulus pupil diameter*))^−1^. All regressors were standardized by computing their z-score.

Trial-based analysis using GLMM indicated that categorization errors were associated with older age and greater pre-stimulus pupil diameter, as revealed by significant effects of age (*p* < 0.001), and pre-stimulus baseline diameter (*p* = 0.010) on task accuracy. The interaction between age and pupil diameter was not significant (*p* = 0.409), suggesting that prestimulus pupil diameter influenced categorization accuracy equally in both groups. Fixed effects accounted for 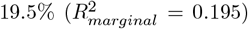 of the total variance observed, and the inclusion of random effects increased the explained variance to 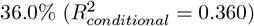. Predicted probability of correct categorization as a function of age and pre-stimulus pupillary diameter is shown in Fig. 3C. A slightly better model 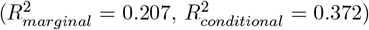 to predict categorization accuracy was obtained by adding stimulus congruence (*p* = 0.002), foreground object type (*p* < 0.001), and their interaction to the previous model (*p* = 0.070). All variables except interactions significantly explained categorization success. More precisely, congruent stimuli and furniture-containing scenes were both associated with better accuracy. Standardized regression estimates, associated p-values, and goodness-of-fit criteria are given in supplementary table S1.

Subject-based TEPRs analysis showed that categorization errors induced greater pupil dilation in older adults (*n* = 20, mean t = 3.013, mean Cohen’s *d* = 0.713, TFCE-corrected; Fig. 3B). This effect could not be observed in the younger group, since most participants’ accuracy was over 85 % (except for one individual), with too few error trials to compute an error-specific median TEPR. These findings were supported by the trial-based GLMM results 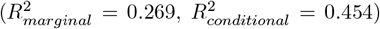, which identified negative effects of age (*p* < 0.001) and dilation amplitude (*p* < 0.001) on categorization success. The non-significant interaction between age and dilation amplitude suggests that this relationship holds consistently across both age groups (*p* = 0.849; table S2).

### 3.3. Pupil dynamics at encoding and subsequent memory performance

Global scene recognition performance at 24h was worse in older participants (mean ± SD = 58.6 ± 6.2 %) than in younger ones (64.8 ± 6.2 %; *t*(89) = 4.803, *p* < 0.001, Cohen’s *d* = 1.007; Fig. 4A). Notably, there were 40 younger and 21 older participants (respectively 89% and 46% of the samples) with overall recognition performance above chance level (≥ 59%). These individuals were included in the analysis relating pupil response at encoding to subsequent stimulus recognition.

**Figure 4.**
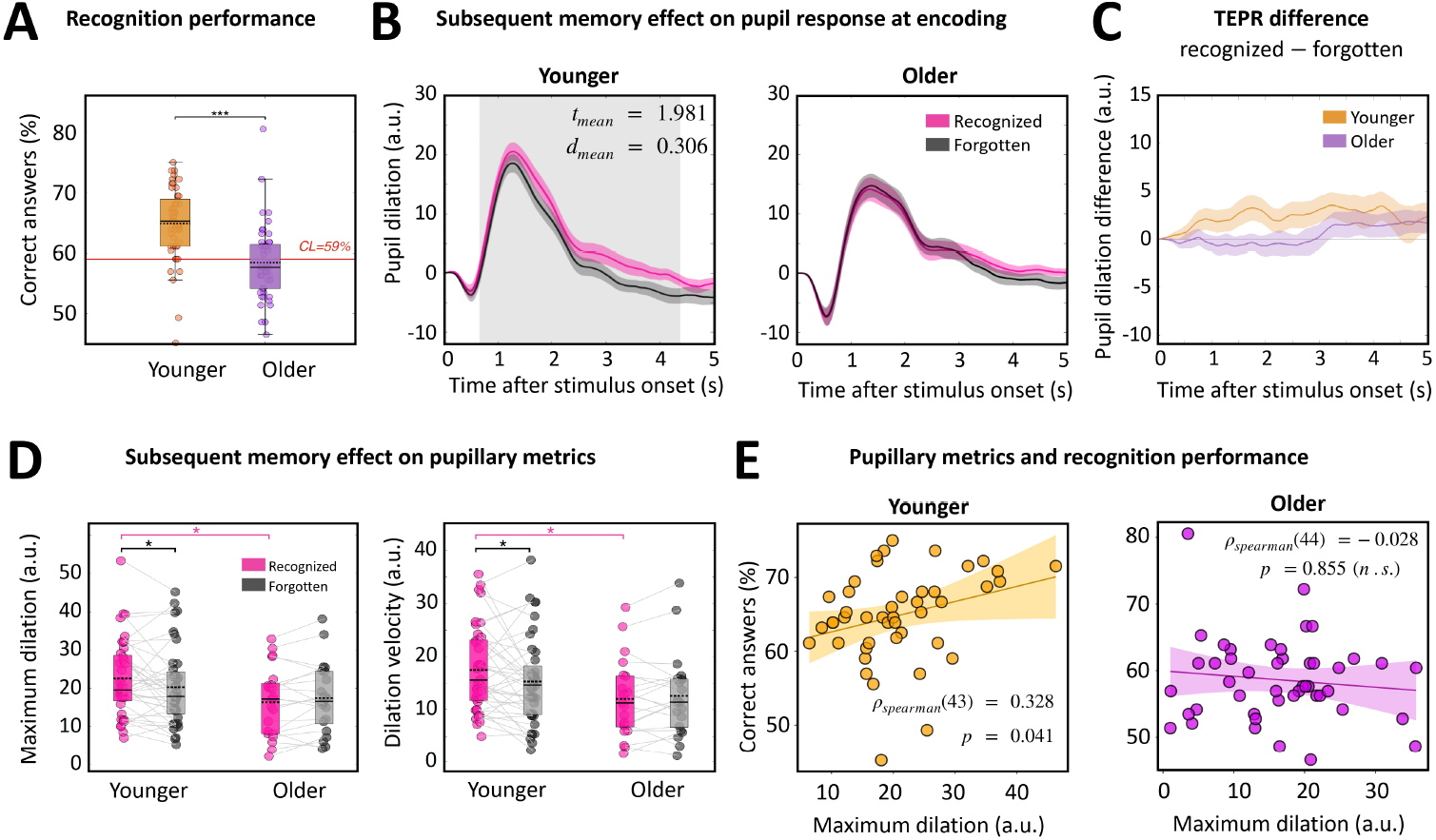
Recognition test (**A**) Recognition performance in younger and older individuals. The red line indicates chance level performance (*<* 59%). (**B**) TEPRs in younger (*n* = 40, left panel) and older (*n* = 21, right panel) individuals, for subsequently recognized and forgotten scenes. Shaded gray area represents the cluster that survived TFCE correction. One-tailed paired t-tests with *H*_1_ : recognized *>* forgotten. Error bar represents s.e.m. (**C**) TEPRs difference (recognized−forgotten) between age groups. No cluster survived TFCE correction. One-tailed independent t-tests with *H*_1_ : younger *>* older. (**D**) Maximum dilation (left) and dilation velocity (right) in both age groups for subsequently recognized and forgotten scenes. (^∗^*p* < 0.05;^∗∗^ *p* < 0.01;^∗∗∗^ *p* < 0.001, FDR correction). The solid and dotted lines in boxplots represent group median and mean values, respectively. (**E**) Recognition performance as a function of maximum dilation in younger (left) and older adults (right).

Younger individuals’ pupil dilation was greater for subsequently recognized relative to forgotten scenes (*n* = 40, mean t = 1.981, mean Cohen’s *d* = 0.306, TFCE-corrected; Fig. 4B, left panel). Greater maximum dilation amplitude was observed for successfully recognized stimuli (*t*(59) = 2.236, *p* = 0.044, Cohen’s *d* = 0.229, one-tailed post-hoc test; Fig. 4D, left panel), as well as faster dilation velocity (*t*(59) = 2.317, *p* = 0.027, Cohen’s *d* = 0.273, one-tailed posthoc test; Fig. 4D, right panel) and greater mean dilation (*t*(59) = 2.625, *p* = 0.033, Cohen’s *d* = 0.318, one-tailed post-hoc test). There were no differences in older participants’ pupillary responses between subsequently recognized and forgotten stimuli (*n* = 21; Fig. 4B, right panel). Finally, when examining the difference in pupillary responses based on memory outcome (*recognized* minus *forgotten*) across age groups, no significant differences were found (Fig. 4C), even when responses were expressed as relative dilation (% of pre-stimulus diameter).

In the younger group, two phasic pupillary metrics (averaged over all trials) positively correlated with global recognition performance, namely dilation amplitude (*ρ*(43) = 0.328, *p* = 0.041, Fig. 4E, left panel) and dilation velocity (*ρ*(43) = 0.359, *p* = 0.041). In older individuals, there were no correlations between phasic pupillary metrics and recognition performance (for dilation amplitude, see Fig. 4E, right panel). To further explore memory effects in the aged group, we compared overall pupillary responses between older subjects who performed at chance level (*n* = 25) and those who performed above chance level (*n* = 21). Following stimulus presentation, we noted a steeper reflexive constriction in subjects who performed above chance level, but this effect did not survive TFCE correction (Fig. S2A), even when considering relative dilation (% of pre-stimulus diameter). In terms of singular phasic metrics, there were no significant differences between groups. Moreover, there was no difference in pre-stimulus diameter (*t*(44) = 1.753, *p* = 0.087, Cohen’s *d* = 0.519; Fig. S2B). Given that the TEPRs showed distinct constriction reflexes depending on recognition accuracy, we further examined the relationship between maximum constriction amplitude and memory performance. This correlation was significant only in the older group (older: *ρ*(44) = −0.413, *p* = 0.009; younger: *ρ*(43) = 0.204, *p* = 0.179; Fig. S2C), indicating that better performers exhibited a steeper constriction reflex.

Finally, the trial-based analysis did not reveal a clear relationship between pupillary metrics and recognition success, as indicated by low goodness-of-fit values of the models (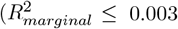; tables S3 & S4). None of the fixed effects, i.e., pre-stimulus diameter and maximum dilation, significantly accounted for the variance in recognition performance. Although the inclusion of congruence and foreground object type as fixed factors did not meaningfully enhance model fit 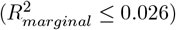, both factors remained statistically significant across models, with improved memory for incongruent and animal-containing stimuli.

### 3.4. Pupil dynamics at encoding and subsequent memory strength

Confidence indices were significantly reduced in the older relative to the younger group (*F* (1, 89) = 23.07, *p* < 0.001) and varied significantly across confidence levels (R vs. K vs. G, *F* (2, 178) = 27.48, *p* < 0.001). No interaction was found between age and confidence level (*p* = 0.848). We indeed observed greater index values for recollection (*t*(89) = 2.163, *p* = 0.041, Cohen’s *d* = 0.531; Fig. 5A), familiarity-based recognition (*t*(89) = 3.591, *p* = 0.001, Cohen’s *d* = 0.634), and guessing (*t*(89) = 2.282, *p* = 0.036, Cohen’s *d* = 0.459) in younger compared to older adults. Furthermore, in younger participants, recollection memory more effectively discriminated old from novel stimuli, compared to familiarity-based memory (*t*(89) = 3.423, *p* = 0.002, Cohen’s *d* = 0.710), which was in turn more efficient than guessing (*t*(89) = 2.254, *p* = 0.036, Cohen’s *d* = 0.436). In older subjects, recollection was also more efficient than familiarity-based recognition (*t*(89) = 3.963, *p* < 0.001, Cohen’s *d* = 0.813), and familiarity-based memory was as accurate as guessing (*p* = 0.204).

**Figure 5.**
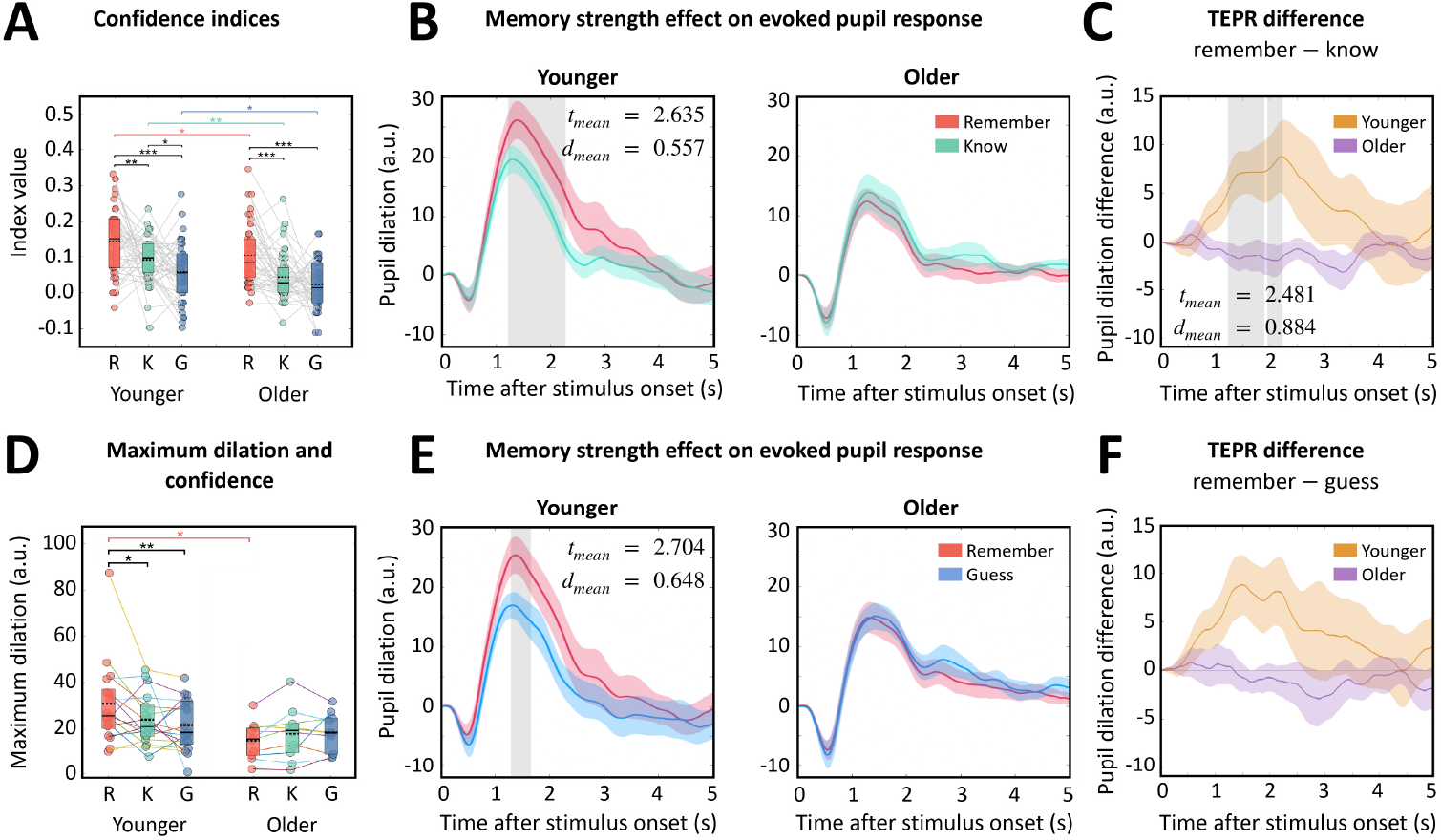
Recognition test (**A**) Recollection (R), familiarity-based memory (K), and guessing (G) confidence indices in both age groups. (**B**) TEPRs in younger (*n* = 20, left panel) and older (*n* = 13, right panel) individuals, for subsequent R and K responses. Shaded gray area represents the cluster that survived TFCE correction. One-tailed paired t-tests with *H*_1_ : remember *>* know. Error bar represents s.e.m. (**C**) TEPRs difference (remember−know) in both age groups. One-tailed independent t-tests with *H*_1_ : younger *>* older. (**D**) Peak dilation at encoding for subsequent R, K and G responses, in both age groups. (**E**) TEPRs in younger (*n* = 21, left panel) and older (*n* = 13, right panel) individuals, for subsequent R and G responses. One-tailed paired t-tests with *H*_1_ : remember *>* guess. (**F**) TEPRs difference (remember − guess) between age groups. One-tailed independent t-tests with *H*_1_ : younger *>* older. (^∗^*p* < 0.05;^∗∗^ *p* < 0.01;^∗∗∗^ *p* < 0.001, FDR correction). The solid and dotted lines in boxplots represent group median and mean values, respectively.

Pupil response analysis as a function of subsequent recognition strength was conducted on 21 younger and 13 older individuals, who showed a sufficient number of valid trials for each experimental condition (R vs. K vs. G). Younger individuals evidenced increased pupil dilation for subsequently recollected scenes, relative to those recognized on a basis of familiarity (*n* = 20, mean t = 2.635, mean Cohen’s *d* = 0.557, TFCE-corrected; Fig. 5B, left panel). Peak dilation amplitude was also significantly higher for recollected stimuli (*t*(31) = 2.670, *p* = 0.018, Cohen’s *d* = 0.542, one-tailed post-hoc test; Fig. 5D). In contrast, no effect of recollection over familiarity-guided recognition was observed in older individuals’ pupillary responses (*n* = 13; Fig. 5B, right panel). Within-subject TEPRs difference between the two conditions (*remember* minus *know*) was greater in the young relative to the older group (mean t = 2.481, mean Cohen’s *d* = 0.884, TFCE-corrected; Fig. 5C). Note that this result remained significant when considering measures of relative dilation (in % of pre-stimulus diameter). Moreover, in younger individuals only, pupil responses to subsequently recollected stimuli showed greater dilation, when compared to those recognized by guessing (*n* = 21, mean t = 2.704, mean Cohen’s *d* = 0.648, TFCE-corrected; Fig. 5E, left panel). An increase in peak dilation amplitude was also observed (*t*(31) = 3.946, *p* = 0.003, Cohen’s *d* = 0.590, one-tailed post-hoc test; Fig. 5D). No effect of recollection over guessing was observed in older participants’ pupil responses (*n* = 13; Fig. 5E, right panel). In addition, when examining the difference in pupillary responses between the two conditions (*remember* minus *guess*) across age groups, the original cluster did not survive TFCE correction (Fig. 5F) even when responses were expressed as relative dilation (% of pre-stimulus diameter). Finally, pupil responses were equivalent for scenes subsequently recognized through familiarity and guessing, in both younger and older groups, and no differences in pupillary dilation metrics were observed.

### 3.5. Effects of stimulus properties on behavior and pupil dynamics: congruence and foreground object type

In both younger and older individuals, object categorization accuracy was impaired for incongruent relative to congruent scenes (*F* (1, 88) = 11.24, *p* = 0.001; post-hoc tests: younger group, *t*(88) = 2.081, *p* = 0.040, Cohen’s *d* = 0.172; older group: *t*(88) = 2.666, *p* = 0.011, Cohen’s *d* = 0.216). Mean RTs for correct trials were longer in response to incongruent scenes (*F* (1, 88) = 18.11, *p* < 0.001) and this effect was exclusively observed in the older group (*Age* × *Congruence* interaction, *F* (1, 88) = 7.08, *p* = 0.009; post-hoc test in the older group: *t*(88) = 4.945, *p* < 0.001, Cohen’s *d* = 0.283). Results are shown on Fig. S3A. In younger individuals only, recognition at 24h was higher for incongruent relative to congruent scenes (*Age* × *Congruence* interaction, *F* (1, 89) = 10.41, *p* = 0.002; post-hoc test in the younger group: *t*(89) = 2.667, *p* = 0.014, Cohen’s *d* = 0.410). Results are shown on Fig. S3B.

An impact of visual incongruence on younger individuals’ TEPRs was observed (*n* = 45, mean t = 2.801, mean Cohen’s *d* = 0.427; Fig. S3C, left panel), resulting in a slightly longer response to incongruent scenes. No difference across congruence conditions was seen in older individuals’ TEPRs (*n* = 46; Fig. S3C, right panel). Overall, there were no differences in any of the phasic pupillary metrics between the two congruence conditions in either group (*p >* 0.163). Furthermore, when comparing differences in pupil time-series between congruence conditions (*congruent* minus *incongruent*) across age groups, no cluster was observed (Fig. S3D).

The foreground object type influenced categorization accuracy (*F* (1, 88) = 4.06, *p* = 0.047), with increased errors in response to animals relative to pieces of furniture in older individuals only (*t*(88) = 2.410, *p* = 0.022, Cohen’s *d* = 0.264; Fig. S4A, left panel). Moreover, shorter RTs were observed for animals relative to pieces of furniture (*F* (1, 88) = 48.08, *p* < 0.001). This was evidenced both in younger (*t*(88) = 2.829, *p* = 0.009) and older (*t*(88) = 7.024, *p* < 0.001) groups. The *Age* × *Object type* interaction was significant (*F* (1, 88) = 8.34, *p* < 0.005; Fig. S4A, right panel). Better recognition performance was evidenced for animal objects relative to pieces of furniture (*F* (1, 89) = 16.71, *p* < 0.001), in both younger (*t*(89) = 3.518, *p* = 0.001) and older groups (*t*(89) = 2.257, *p* = 0.032). There was no significant *Age* × *Object type* interaction on recognition performance (*p* = 0.363; Fig. S4B).

Based on subject-based TEPRs analysis, there were no significant differences between pupil dilation patterns in response to animals and pieces of furniture, in any age groups (Fig. S4C). Note that in younger individuals, pupil dilation was increased in response to animals, although no clusters survived TFCE correction. No significant differences in any phasic pupillary metrics were observed between the two object types in either group (*p >* 0.093). Moreover, when comparing the differences in pupil dilation between both object types (*animal* minus *furniture*) across age groups, no significant clusters were observed (Fig. S4D).

Finally, the trial-based analysis showed that stimulus properties poorly explained pupillary dilation (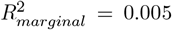 and 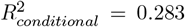). However, the foreground object type (*p* = 0.003) and its interaction with scene congruence (*p* < 0.001) both emerged as significant fixed effects. Dilation amplitude was greater in response to animals than to furniture, especially when the animals appeared in incongruent backgrounds (supplementary table S5).

## 4. Discussion

This study aimed to investigate the relationship between pupil characteristics, both at rest and task-related, and behavioral performance in healthy younger and older adults. Participants performed fixation and visual scene categorization tasks, and were further tested on long-term memory of implicitly encoded scenes. Our results indicate a global age-related reduction in resting-state pupil diameter, and in phasic pupil dilation amplitude and velocity during the categorization task. We also demonstrated a link between pupil characteristics and cognitive performance that evolved in older age. Our data are in line with previous results reporting decreased basal pupillary diameter in older adults (Tekin et al., 2018), independently of luminance (Guillon et al., 2016) or light wavelength conditions (Lobato-Rincan et al., 2014), as well as lower pupil phasic response amplitude during a variety of cognitive tasks (Riley et al., 2024; He et al., 2020; Hammerer et al., 2017; Van Gerven et al., 2004). Age-related changes in both basal and evoked pupillary properties are considered to reflect altered central LC-driven mechanisms, rather than purely mechanical defects of iris muscles (Mather and Harley, 2016; Korczyn et al., 1976). Continuous inhibitory modulation exerted by the LC on the parasympathetic constriction reflex loop (Joshi and Gold, 2020; Szabadi, 2018) may become less effective with age, contributing to a global decrease in pupil size (Huang et al., 2024; Bittner et al., 2014). In addition, changes in arousal dynamics with age, resulting from modified tonic and phasic LC activity (Weinshenker, 2018), impact pupil diameter time-course both at rest and during tasks. Therefore, the alterations of pupil dynamics in older subjects, observed in the present study and by others, are likely a reliable proxy of the well-documented age-related changes reported in the LC in post-mortem (Ehrenberg et al., 2018; Theofilas et al., 2017; Braak and Del Tredici, 2011) and neuroimaging (Beardmore et al., 2024; Holland et al., 2021; Betts et al., 2017) studies.

During the fixation task, besides the diminution of mean pupil diameter, older age was associated with a decrease in the amplitude of resting-state spontaneous pupil fluctuations. To our knowledge, such age-related changes in basal pupil dynamics at rest have not been reported previously. In young individuals, it has been proposed that pupil variability at rest reflects spontaneous shifts between different mental states due to self-regulation of arousal (Aminihajibashi et al., 2019). In our experiment, pupil size fluctuations in young subjects could denote self-initiated attentional focus on the fixation task after periods of mind-wandering, as previously observed (Grandchamp et al., 2014). These shifts between mental states would be driven by variations in tonic LC discharges, and would influence activity and connectivity in default-mode and executive control brain networks (Wu et al., 2025; Yellin et al., 2015; Eldar et al., 2013). Our data show smaller pupil fluctuations in the older group, consistent with a decrease in the frequency of mind-wandering as a function of age that has been reported across a variety of experimental paradigms (for review see Maillet and Schacter (2016)). Interestingly, a recent study reported lower pre-stimulus pupil variability in older compared to young adults, during a visual change detection task (Moran et al., 2025). In this study, subjects reported their mental state following change detection in each trial, and more focused task engagement was observed in older adults. Our results on the fixation task may therefore indicate that older participants had reduced mind-wandering during rest, which could be captured using pupil measures.

In the rapid scene categorization task, global performance was overall high in the two groups. It was diminished in older versus younger individuals, as previously reported (Rémy et al., 2013). In both age groups, trial-based analyses showed that categorization accuracy was related to pupil phasic dilation amplitude and pre-stimulus diameter. Firstly, phasic pupillary dilation was greater for scenes that were wrongly categorized. Likely, this dilation was a marker of error-driven transient arousal, as previously observed using a fast Stroop paradigm (Compton et al., 2021), and therefore did not condition trial performance. Secondly, we found that pre-stimulus pupil size predicted subsequent categorization performance, with a larger pre-stimulus pupil diameter associated with scene miscategorization. Pre-stimulus pupil size was likely related to the amount of attention towards the task (Irons et al., 2017). Indeed, moment-to-moment fluctuations in pupil diameter capture temporal changes in arousal levels and varying vigilance states (McGinley et al., 2015). Using a simple visual detection task that required sustained attention, Unsworth and Robison (2016) observed that a larger pre-stimulus pupil diameter was linked to increased reaction time. In the latter study, periodical thought probes were presented during the task, and greater pre-stimulus pupil size was associated with participants’ self-reports of external distraction, inducing delayed responses. Consistently, such tonic pupil spontaneous dilations have been associated with activation of the salience network, involved in orientation to salient stimuli (Schneider et al., 2016). Our data may thus indicate that a larger baseline pupil diameter reflected heightened arousal and increased distractibility. Stimulus processing was less optimal, thereby increasing the likelihood of categorization errors. This interpretation is in agreement with the adaptive gain theory of the LC-NA system function (Aston-Jones and Cohen, 2005). Within this framework, elevated tonic LC activity enables exploration for alternative behaviors in a changing environment, i.e., external distraction, with disengagement from the current task. Conversely, moderate levels of tonic LC activity allow for optimal task-relevant processing. With regard to aging, in our study, although pre-stimulus pupil size was globally reduced in older subjects, its variability conditioned task performance in a similar way to younger subjects, so that, within a trial, larger baseline pupil diameter was related to categorization error. Therefore, higher proneness to distraction with age could have contributed to more frequent errors. A previous functional MRI study (Lee et al., 2020) reported that age-related changes in resting-state connectivity between the LC and the salience network may account for failure to prioritize task-relevant inputs and greater distractibility. Our data suggest that, in addition, non-optimal elevated levels of tonic LC activity during task performance could be more frequent in older age, and possibly enhance vulnerability to distraction.

Our study revealed a link between phasic pupil response and long-term recognition performance in younger participants. Within-subjects TEPR analyses showed that greater dilation at encoding was related to successful scene recognition 24 hours later. Phasic dilation was also greater for stimuli subsequently recollected, compared to those recognized with a sense of familiarity or by guessing. Our data in young subjects suggest that levels of phasic arousal, likely reflecting attentional focus and cognitive effort for rapid object categorization, influenced incidental encoding processes and storage in long-term memory. This is in agreement with previous studies that showed, using intentional word encoding (Pitem and Mama, 2025; Miller et al., 2019; Bergt et al., 2018; Kucewicz et al., 2018; Papesh et al., 2012) or incidental scene encoding (Pilarczyk et al., 2022) tasks, better memory for stimuli that elicited higher arousal at encoding. Conversely, other studies reported stronger pupil constriction during incidental encoding of visual objects (Kafkas and Montaldi, 2011) or scenes (Naber et al., 2013) that were subsequently recognized with high confidence. The latter two studies used passive encoding tasks, with free visual exploration. Notably, Kafkas and Montaldi (2011) showed that more active object exploration, i.e. higher number and shorter duration of gaze fixations, also promoted object memorization, possibly thanks to deeper encoding of visual details and independently of global arousal levels. In our experiment, brief stimulus presentation prevented eye movements and scene exploration. Also, the rapid object categorization task required focused attention and active cognitive processing. This may explain why we observed phasic pupil responses which were of higher amplitude relative to those elicited by passive viewing (Kafkas and Montaldi, 2011; Naber et al., 2013), and more similar to those reported during intentional encoding (Pitem and Mama, 2025; Miller et al., 2019; Bergt et al., 2018; Kucewicz et al., 2018; Papesh et al., 2012). Our results are consistent with a previous study combining pupil and fMRI measures (Clewett et al., 2018), which reported that under threat-related arousal, higher pupil dilation at encoding predicted better memory for prioritized visual scenes. Importantly, in the latter study, pupil dilation was accompanied by an increase in LC activity, which was thought to promote narrowed attentional focus and efficient neural processing in scene-selective regions of the ventral pathway. Accordingly, our pupil results in the young group may indicate that, even without any emotionally-arousing context, some scenes elicited greater phasic arousal and captured higher attentional and cognitive resources at encoding (Miller et al., 2019). Concomitant phasic LC activity and NA release may have enhanced neural gain in the executive control network and ventral visual cortex (Clewett et al., 2018; Eldar et al., 2013). Since we found a higher pupil response for scenes that were subsequently recollected, it is also possible that phasic co-release of both noradrenaline and dopamine in the hippocampus (Dahl et al., 2023) through direct projections from the LC (Sara, 2009; Samuels and Szabadi, 2008a,b) promoted synaptic plasticity. Indeed, the formation of stronger hippocampal representations could have enabled easier contextual retrieval and higher recollection success, as observed in our young group.

In older participants, a drop in scene recognition performance was observed, and affected both recollection- and familiarity-guided processes. This is consistent with previous reports of agerelated decline in recollection performance, and to a lesser degree in familiarity-based recognition (Koen and Yonelinas (2014) for review). Moreover, there was no relationship between phasic pupil response at scene encoding and subsequent scene memory in older adults. Therefore, while scene encoding strength was promoted by higher phasic arousal in the young group, no such benefit on memory encoding was evidenced in the older group. This may reveal diminished involvement of the LC-NA system during scene processing in the older group, which could have partly accounted for the decline in recognition performance. Notably, in our experiment, trial-to-trial variability in phasic pupil response amplitudes was significantly reduced in older age. Scene properties, such as incongruence or the presence of animals, did not impact transient pupil dilation, suggesting less bottom-up influence of visual salient information on LC activity. In agreement, during scene recognition, older adults evidenced smaller pupil phasic dilation in response to emotional versus neutral novel scenes, relative to younger adults (Hammerer et al., 2017). Therefore, age-related reduction of arousal at encoding may result in decreased NA release and neural gain, inducing altered activity in regions involved in executive control and visual processing, in line with previous results combining pupil and fMRI measures in older subjects (Lee et al., 2018). Also consistent with our observations, neuroimaging studies in older healthy individuals have demonstrated that poorer long-term memory performance was associated with reduced structural integrity in the LC (Elman et al., 2021; Liu et al., 2020; Hammerer et al., 2018), reduced LC activity (Prokopiou et al., 2022), and decreased restingstate connectivity between the LC and ventral visual cortex (Jacobs et al., 2015). In particular, the rostral region of the LC is the first to selectively show structural and functional changes with age, while the caudal part may stay better preserved (Veréb et al., 2023; Dahl et al., 2019; Betts et al., 2017). Greater structural integrity in the rostral LC has been linked to better episodic memory performance (Dahl et al., 2019), and higher resting-state functional connectivity between the rostral LC and prefrontal and temporal regions has been associated with higher emotional memory scores (Veréb et al., 2023). Therefore, during phasic LC activation in response to stimuli, age-related deterioration of rostral LC projections to the forebrain could contribute to the decline in memory encoding, while having little impact on pupil dilation, which is controlled through caudal LC projections to the spinal cord and the brainstem (Szabadi, 2018; Samuels and Szabadi, 2008a,b). Consistently, under threat-related arousal, a lack of correlation between pupil dilation and cortical fMRI activity has been reported specifically in older adults (Lee et al., 2018). In our study, such age-related LC changes may account for the lack of relationship between pupil dilation amplitude and recognition performance in the older group.

There is a large body of evidence showing that aggregates of hyper-phosphorylated tau, i.e. pre-tangles, occur initially in LC neurons, where they gradually accumulate throughout the adult span and further propagate to brain regions (Harley et al., 2021; Theofilas et al., 2017; Braak et al., 2011; Braak and Del Tredici, 2011). Additionally, LC integrity and activity, assessed in vivo with MRI, have been both related to cerebral markers for Alzheimer’s disease (AD) (Prokopiou et al., 2022; Jacobs et al., 2021). This suggests that pupil dynamics could be a relevant marker to help identify subjects at risk for cognitive impairment and AD. Locus coeruleus neuronal loss as a consequence of tau aggregates (Theofilas et al., 2017), and concomitant decline in LC connectivity (Jacobs et al., 2015) and noradrenergic signaling (Portela Moreira et al., 2023; Pan et al., 2020), are thought to occur at prodromal Braak stages III-IV of AD. Consistently, diminished baseline pupil diameter (Fotiou et al., 2009; Prettyman et al., 1997) and smaller phasic pupillary response (Haj et al., 2022; Granholm et al., 2017; Dragan et al., 2017) have been reported in AD patients. Preceding the symptomatic phase, the occurrence of intra-neuronal pre-tangles could contribute to tonic and phasic LC hyperactivity, and increased cerebral NA levels (Weinshenker, 2018). In asymptomatic individuals, tonic and phasic pupil measures may thus depend on tau burden in LC neurons. Our results on tonic pre-stimulus pupil measures are coherent with an age-related tonic LC hyperactivity, which would impact sustained engagement on task, increase distractibility, and deteriorate task performance (Aston-Jones and Cohen, 2005). Thus, baseline pupil measures during cognitive tasks may represent a useful marker of tonic LC hyperactivity. Note that, during rest, tonic LC activity could be difficult to evaluate through pupillary measures, since resting-state pupil diameter may be confounded by LC influence on the parasympathetic pupil reflex (Huang et al., 2024) and may strongly depend on fluctuations in mental states that evolve with age (Moran et al., 2021). Regarding phasic measures, we found that pupil response amplitude was globally reduced in the older group, which contradicts the previously hypothesized phasic LC hyperactivity with age (Weinshenker, 2018). A possibility is that greater phasic pupil responses may be measured in middle-aged individuals, but no more in older age (Riley et al., 2024). Interestingly, larger phasic pupillary dilation has been linked to polygenic risk for AD in cognitively-healthy middle-aged adults (Kremen et al., 2019). In line with this, phasic LC activity in a transgenic rat model of AD producing endogenous tau was elevated when tau pathology was confined to the LC, but reduced at more severe stages (Kelberman et al., 2023). Consequently, evoked pupil measures during cognitive tasks may be mostly relevant in middle-aged individuals as a marker of LC phasic hyperactivity. Finally, our data indicate that the parasympathetic pupil light reflex (PLR) could constitute an additional pupillary marker of interest. When exploring determinants of TEPRs within the older group, we found that steeper reflexive constriction amplitude was associated with higher recognition performance. The PLR amplitude is known to reduce with age, and further diminish in individuals with mild cognitive impairment and AD (Chougule et al., 2019). Moreover, PLR amplitude has been positively related to MMSE scores (Bittner et al., 2014) and general cognitive abilities (Chen et al., 2022). As the LC exerts inhibitory modulation of the parasympathetic reflex, tau aggregates in LC neurons may influence PLR amplitude. Tau pathology within the EWN, which is part of the reflex loop, may also be a major cause for age-related changes in PLR constriction (Scinto et al., 2001), suggesting that this pupil marker may reflect more global tau burden in the brainstem and be related to cognitive decline. Overall, our results on pupil dynamics in older age are promising and encourage further investigations, combining pupillometry at rest and during tasks, along with neuroimaging assessment of LC integrity and function, and detailed cognitive evaluation. Longitudinal research is mandatory and will help determine whether pupil markers can reliably predict future cognitive decline and AD pathology, thereby supporting their potential as a non-invasive tool to identify at-risk individuals.

Considering our study’s limitations, we note firstly that the memory task was challenging, with fewer than half of our older participants performing above chance on the recognition test. Investigation of memory effects on pupil responses was thus conducted on relatively small groups, although such sample sizes are not uncommon in the cognitive pupillometry literature on the subsequent memory effect (Kucewicz et al., 2018; Hammerer et al., 2017; Papesh et al., 2012; Võ et al., 2008). Our experiment was designed to highlight LC-driven pupil effects and potential age-related changes. The rapid categorization task required sustained attention and active stimulus processing, while permitting incidental encoding. Brief scene presentation prevented eye movements and subsequent pupil foreshortening that may have impaired accuracy in pupil measurement (Mathôt and Vilotijević, 2022). Moreover, it allowed isolating the effects of phasic arousal on subsequent memory, independently of scene exploration. Finally, high task difficulty resulted in large between-subject performance variability in both age groups, which avoided ceiling effects and optimized analysis of correlations with pupil responses. However, the exclusion of many older participants to explore the link between pupil dynamics and memory, and related age effects, remains a limitation in our results, and further replication on larger groups is required. Secondly, we found that scene properties had a significant influence on pupil phasic responses. Stimulus-specific effects are rarely reported in studies employing a diverse range of visual natural stimuli. Although in our study the various stimuli explained only a slight part of the variance in pupil responses, animals in incongruent scenes globally elicited greater pupil dilation, suggesting stimulus-specific effects in our results. Still, the experimental choice for using these stimuli was to optimize potential LC-driven effects that are known to be impaired with age, such as selective attention to salient objects in scenes for rapid categorization, and conflict resolution for processing incongruent scenes (Rémy et al., 2020). This paradigm induced trial-to-trial pupil response variability while testing non-emotional episodic memory, bringing novel and complementary results to previous studies that used emotional scenes (Hammerer et al., 2017) or an emotionally-arousing context of encoding (Clewett et al., 2018; Lee et al., 2018). Nevertheless, replication of our findings using different sets of visual neutral stimuli, with a variety of objects or scene categories, would be informative.

### 5. Conclusion

In summary, our study evidenced age-related changes in pupil dynamics. Older adults exhibited reduced pupil size and variability over time, both at rest and during a visual implicit encoding task. Among young adults, greater transient pupil dilation during stimulus encoding was linked to better recognition and confidence. This association was not observed in older adults, potentially because their pupil responses were more uniform across different trials. Assuming that pupil dynamics reflect moment-to-moment variations in LC discharge and NA release, our results suggest that the benefit provided by the LC-NA system for successful encoding may disappear in older age, and that changes in this system could contribute to age-related initial decline in episodic memory. Our results also highlight the strong potential of pupillometry as a non-invasive, in vivo technique for early detection of LC dysfunction.

## Acronyms and Abbreviations

AD: Alzheimer’s disease
EWN: Edinger-Westphal nucleus
FDR: False discovery rate
GLMM: Generalized linear mixed model
LC: Locus coeruleus
MMSE: Mini-mental state examination
MRI: Magnetic resonance imaging
NA: Noradrenaline
PLR: Pupillary light reflex
RKG: Remember know guess
RT: Response time
TEPR: Task-evoked pupillary response
TFCE: Threshold-free cluster enhancement

## CRediT author statement

**Adrian RUIZ CHIAPELLO**: Writing - Original Draft, Investigation, Methodology, Software, Data curation, Formal analysis, Visualization, Validation **Enzo BUSCATO**: Investigation, Data curation **Muriel MESCAM**: Writing - Review and Editing **Isabelle BERRY**: Funding acquisition **Alexandra PRESSIGOUT**: Resources, Writing - Review and Editing **Andrea ALAMIA**: Conceptualization, Methodology, Writing - Review and Editing **Florence REMY**: Supervision, Funding acquisition, Project administration, Conceptualization, Methodology, Writing - Original Draft, Writing - Review and Editing.

## Acknowledgement

We would like to express our thanks to Loanne Courdeau and Emie Just for their dedication and help with the project. We also thank Emmanuel Barbeau for his advice and exchanges.

## Supplementary material

### Supplementary figures

**Figure S1:**
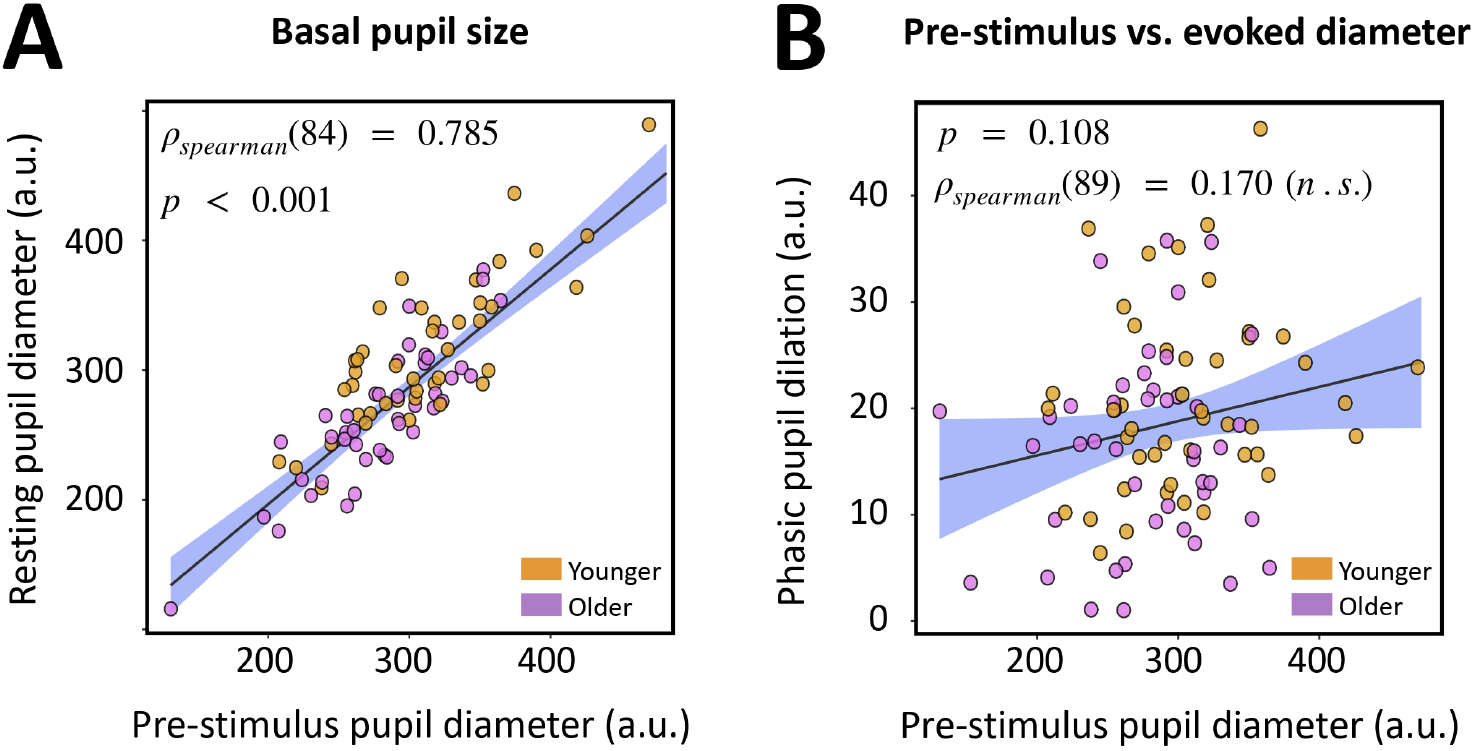
Pre-stimulus pupil diameter (averaged over all trials for each participant) as a function of (**A**) resting-state average pupil diameter, and (**B**) task-evoked pupillary maximum dilation.

**Figure S2:**
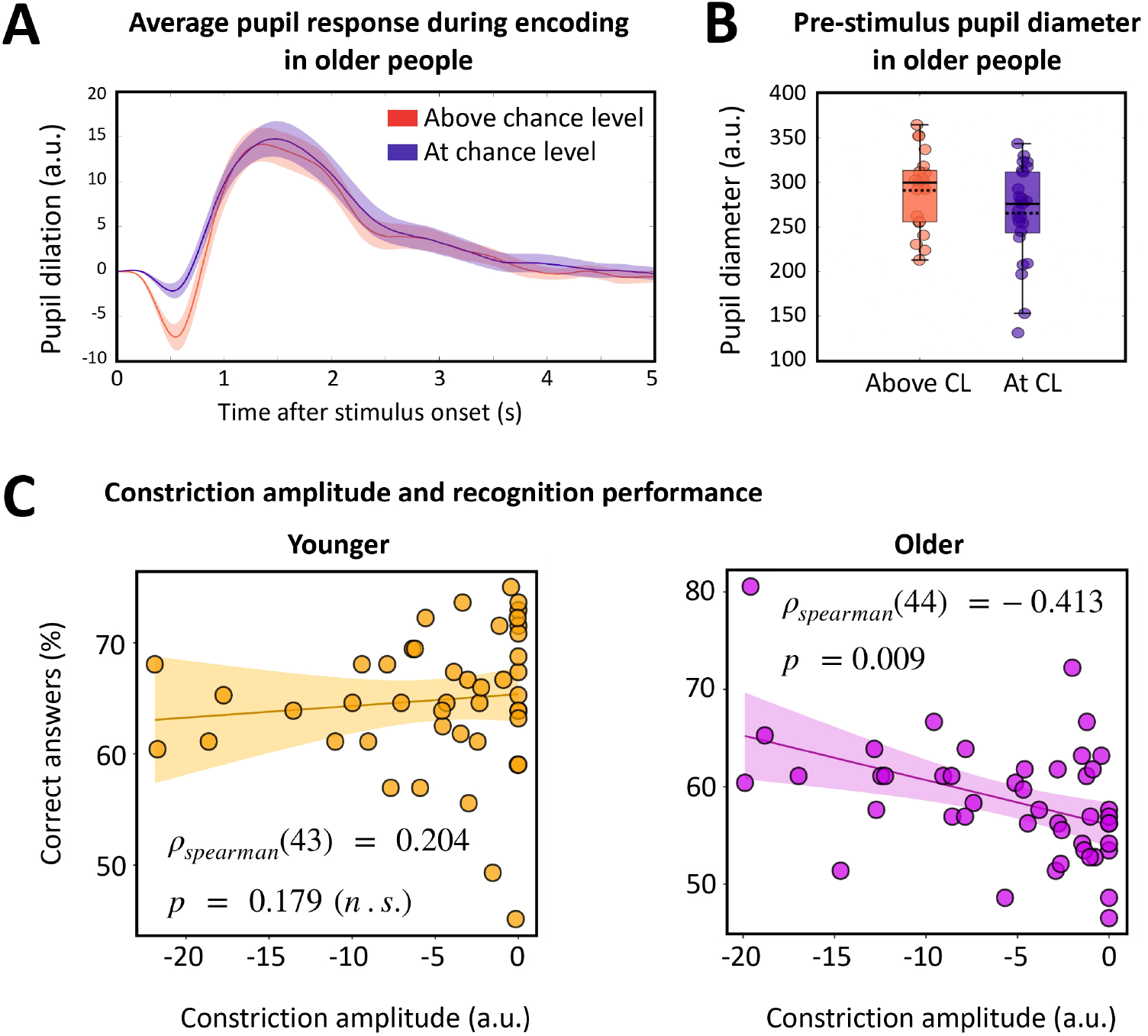
(**A**) TEPRs of older participants above chance level (*n* = 21) and at chance level (*n* = 25) on the recognition test. Two-tailed independent t-tests with *H*_1_ : above chance level *≠* at chance level. (**B**) Median pre-stimulus pupil diameter in older individuals above and at chance level. (**C**) Recognition performance as a function of constriction amplitude in younger (left) and older adults (right). The solid and dotted lines in boxplots represent group median and mean values, respectively.

**Figure S3:**
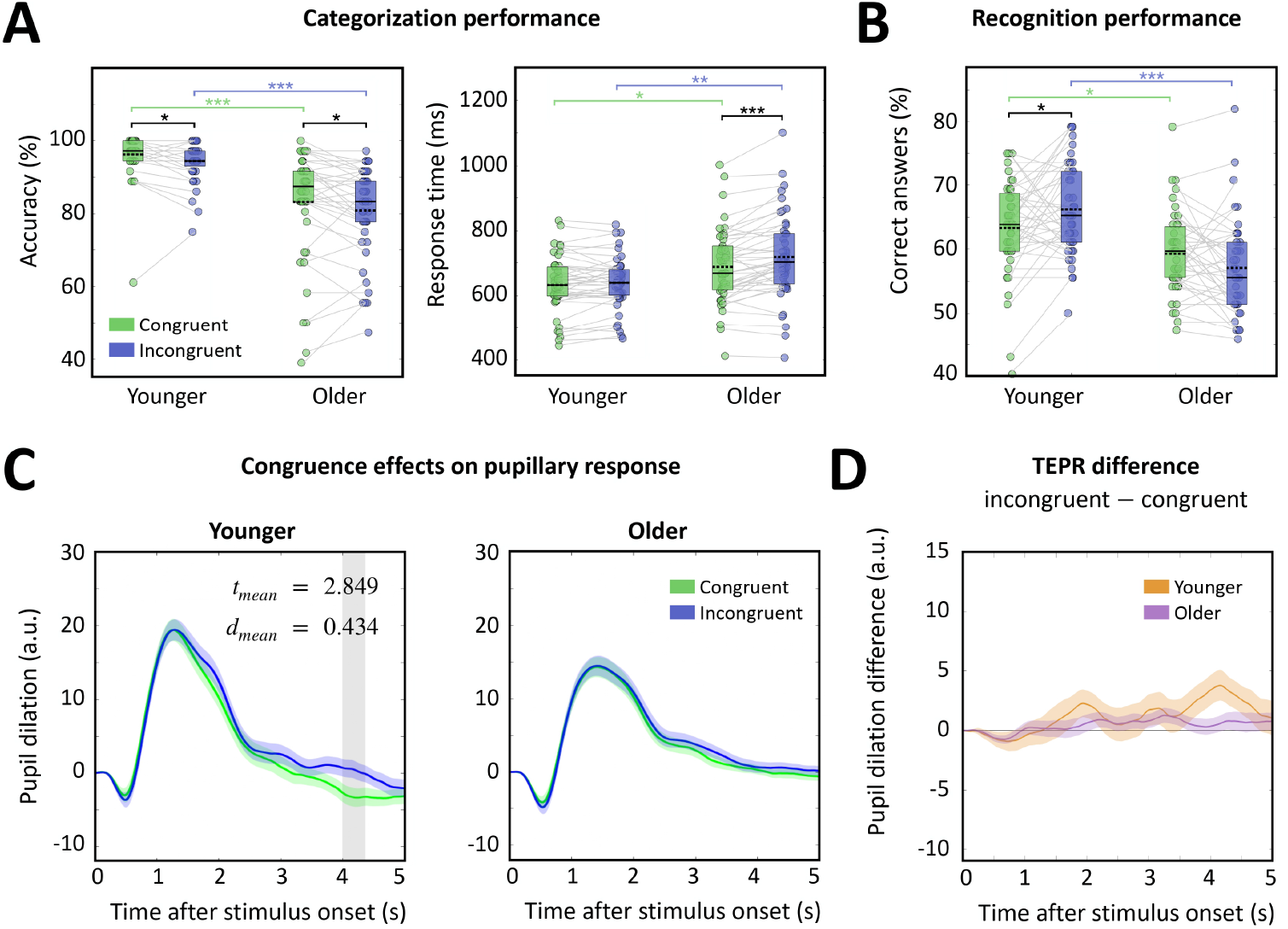
(**A**) Categorization accuracy (left) and mean response time on correct trials (right) for congruent and incongruent scenes in both age groups. (**B**) Recognition performance for congruent and incongruent stimuli in both age groups. (**C**) TEPRs in younger (*n* = 45, left panel) and older (*n* = 46, right panel) individuals, for congruent and incongruent scenes. Shaded gray area represents the cluster that survived TFCE correction. One-tailed paired t-tests with *H*_1_ : incongruent *>* congruent. Error bar represents s.e.m. (**D**) TEPRs difference (incongruent − congruent) in both age groups. One-tailed paired t-tests with *H*_1_ : younger *>* older. (^∗^*p* < 0.05;^∗∗^ *p* < 0.01;^∗∗∗^ *p* < 0.001, FDR correction). The solid and dotted lines in boxplots represent group median and mean values, respectively.

**Figure S4:**
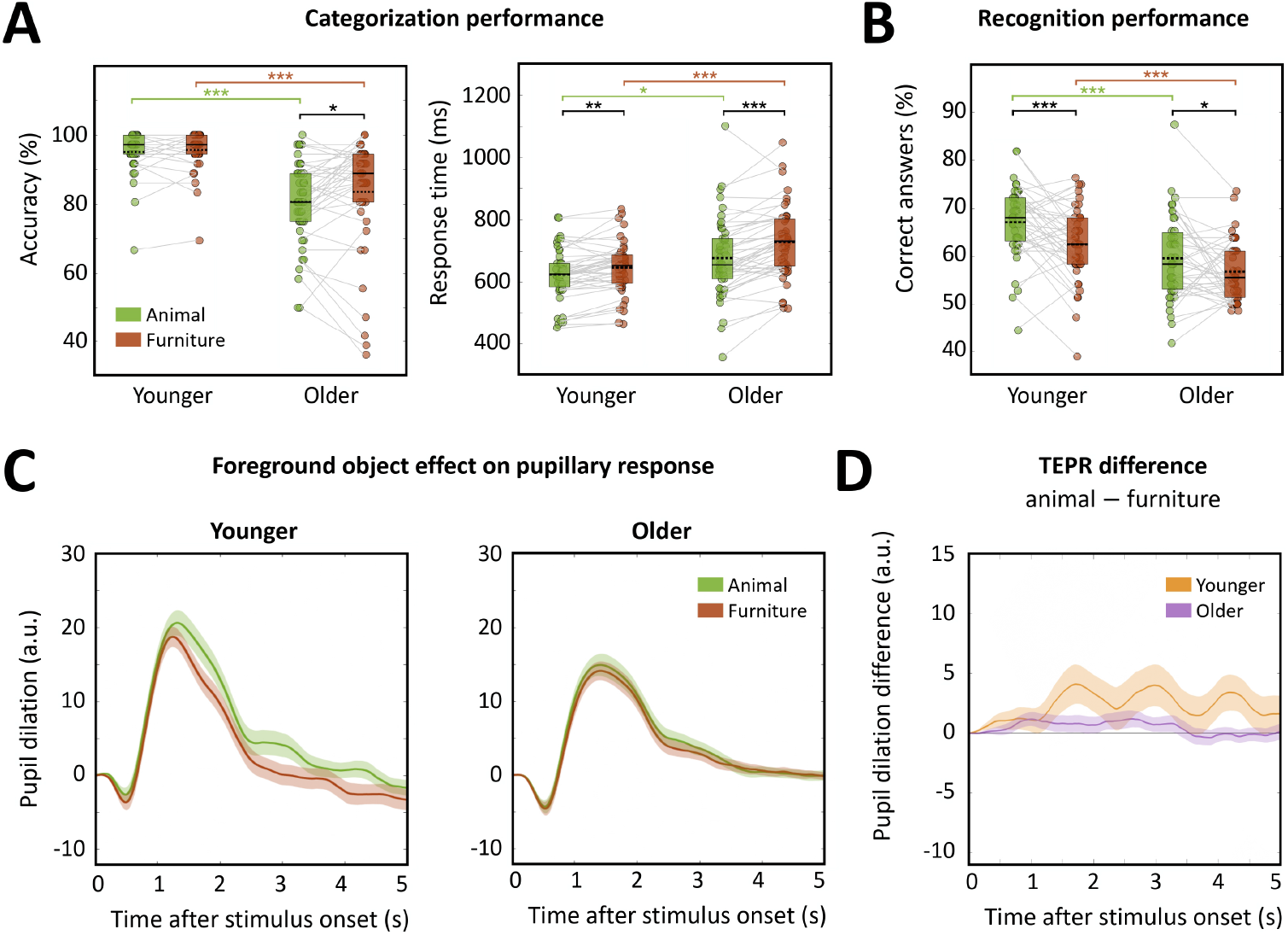
(**A**) Categorization accuracy (left) and mean response time on correct trials (right) for animal and furniture objects in both age groups. (**B**) Recognition performance for animal and furniture objects in both age groups. (**C**) TEPRs in younger (*n* = 45, left panel) and older (*n* = 46, right panel) individuals, for animal and furniture objects. No cluster survived TFCE correction. One-tailed paired t-tests with *H*_1_ : animal *>* object. Error bar represents s.e.m. (**D**) TEPRs difference (animal − object) in both age groups. One-tailed paired t-tests with *H*_1_ : younger *>* older. (^∗^*p* < 0.05;^∗∗^ *p* < 0.01;^∗∗∗^ *p* < 0.001, FDR correction). The solid and dotted lines in boxplots represent group median and mean values, respectively.

### Supplementary tables

**Table S1:**
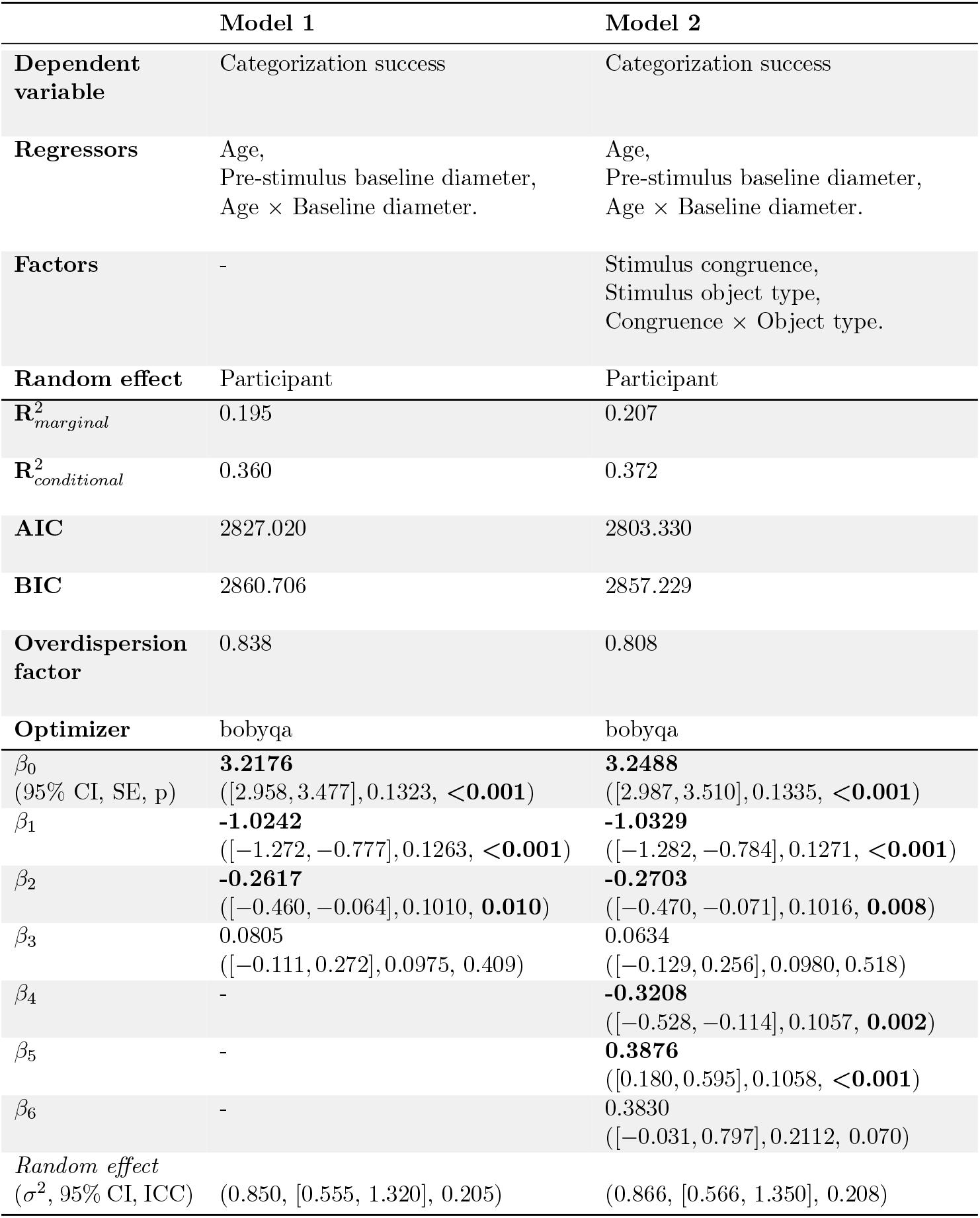
Comparison of model parameters, model diagnostics, regression estimates and associated p-values, between two implemented generalized linear mixed models. **Model 1** is given by the following formula: logit(ℙ[*Categorization success* = 1]) = *β*_0_ + *β*_1_ · *Age* + *β*_2_ · *Pre-stimulus pupil diameter* + *β*_3_ · (*Age* × *Pre-stimulus pupil diameter*) + *Participant*. Formula of **model 2** is: logit(ℙ[*Categorization success* = 1]) = *β*_0_ + *β*_1_ · *Age* + *β*_2_ · *Pre-stimulus pupil diameter* + *β*_3_ · (*Age* × *Pre-stimulus pupil diameter*) + *β*_4_ · *Congruence* + *β*_5_ · *Object type* + *β*_6_ · (*Congruence* × *Object type*) + *Participant*. All regressors were z-scored.

|  | **Model 1** | **Model 2** |
| --- | --- | --- |
| **Dependent variable** | Categorization success | Categorization success |
| **Regressors** | Age,<br>Pre-stimulus baseline diameter,<br>Age × Baseline diameter. | Age,<br>Pre-stimulus baseline diameter,<br>Age × Baseline diameter. |
| **Factors** | - | Stimulus congruence,<br>Stimulus object type,<br>Congruence × Object type. |
| **Random effect** | Participant | Participant |
| $R^2_{\textit{marginal}}$ | 0.195 | 0.207 |
| $R^2_{\textit{conditional}}$ | 0.360 | 0.372 |
| **AIC** | 2827.020 | 2803.330 |
| **BIC** | 2860.706 | 2857.229 |
| **Overdispersion factor** | 0.838 | 0.808 |
| **Optimizer** | bobyqa | bobyqa |
| $\beta_0$<br>(95% CI, SE, p) | **3.2176**<br>([2.958, 3.477], 0.1323, <0.001) | **3.2488**<br>([2.987, 3.510], 0.1335, <0.001) |
| $\beta_1$ | **-1.0242**<br>([-1.272, -0.777], 0.1263, <0.001) | **-1.0329**<br>([-1.282, -0.784], 0.1271, <0.001) |
| $\beta_2$ | **-0.2617**<br>([-0.460, -0.064], 0.1010, **0.010**) | **-0.2703**<br>([-0.470, -0.071], 0.1016, **0.008**) |
| $\beta_3$ | 0.0805<br>([-0.111, 0.272], 0.0975, 0.409) | 0.0634<br>([-0.129, 0.256], 0.0980, 0.518) |
| $\beta_4$ | - | **-0.3208**<br>([-0.528, -0.114], 0.1057, **0.002**) |
| $\beta_5$ | - | **0.3876**<br>([0.180, 0.595], 0.1058, <0.001) |
| $\beta_6$ | - | 0.3830<br>([-0.031, 0.797], 0.2112, 0.070) |
| *Random effect*<br>($\sigma^2$, 95% CI, ICC) | (0.850, [0.555, 1.320], 0.205) | (0.866, [0.566, 1.350], 0.208) |

**Table S2:** Comparison of model parameters, model diagnostics, regression estimates and associated p-values, between two implemented generalized linear mixed models. **Model 1** is given by the following formula: logit(ℙ[*Categorization success* = 1]) = *β*_0_ + *β*_1_ · *Age* + *β*_2_ · *Maximum dilation* + *β*_3_ · (*Age* × *Maximum dilation*) + *Participant*. Formula of **model 2** is: logit(ℙ[*Categorization success* = 1]) = *β*_0_ +*β*_1_ · *Age* +*β*_2_ · *Maximum dilation* + *β*_3_ · (*Age* × *Maximum dilation*) + *β*_4_ · *Congruence* + *β*_5_ · *Object type* + *β*_6_ · (*Congruence* × *Object type*) + *Participant*. All regressors were z-scored.

|  | Model 1 | Model 2 |
| --- | --- | --- |
| **Dependent variable** | Categorization success | Categorization success |
| **Regressors** | Age,<br>Maximum dilation,<br>Age × Maximum dilation. | Age,<br>Maximum dilation,<br>Age × Maximum dilation. |
| **Factors** | - | Stimulus congruence,<br>Stimulus object type,<br>Congruence × Object type. |
| **Random effect** | Participant | Participant |
| $R^2_{\textit{marginal}}$ | 0.261 | 0.269 |
| $R^2_{\textit{conditional}}$ | 0.446 | 0.454 |
| **AIC** | 2367.720 | 2357.370 |
| **BIC** | 2400.562 | 2409.925 |
| **Overdispersion factor** | 0.765 | 0.734 |
| **Optimizer** | bobyqa | bobyqa |
| $\beta_0$<br>(95% CI, SE, p) | **3.3644**<br>([3.051, 3.678], 0.1598, <0.001) | **3.3960**<br>([3.080, 3.712], 0.1611, <0.001) |
| $\beta_1$ | **-1.2298**<br>([-1.529, -0.930], 0.1528, <0.001) | **-1.2340**<br>([-1.535, -0.933], 0.1535, <0.001) |
| $\beta_2$ | **-0.6958**<br>([-0.839, -0.553], 0.0730, <0.001) | **-0.6837**<br>([-0.827, -0.540], 0.0734, <0.001) |
| $\beta_3$ | 0.0041<br>([-0.136, 0.145], 0.0717, 0.954) | 0.0137<br>([-0.128, 0.155], 0.0721, 0.849) |
| $\beta_4$ | - | **-0.2939**<br>([-0.519, -0.068], 0.1151, **0.011**) |
| $\beta_5$ | - | **0.3278**<br>([0.102, 0.553], 0.1150, **0.004**) |
| $\beta_6$ | - | 0.1422<br>([-0.309, 0.593], 0.2301, 0.537) |
| *Random effect*<br>($\sigma^2$, 95% CI, ICC) | (1.10, [0.72, 1.72], 0.251) | (1.11, [0.72, 1.74], 0.252) |

**Table S3:** Comparison of model parameters, model diagnostics, regression estimates and associated pvalues, between two implemented generalized linear mixed models. **Model 1** is given by the following formula: logit(ℙ[*Recognition success* = 1]) = *β*_0_ + *β*_1_ · *Age* + *β*_2_ · *Pre-stimulus pupil diameter* + *β*_3_ · (*Age* × *Pre-stimulus pupil diameter*) + *Participant*. Formula of **model 2** is: logit(ℙ[*Recognition success* = 1]) = *β*_0_ + *β*_1_ · *Age* + *β*_2_ · *Pre-stimulus pupil diameter* + *β*_3_ · (*Age* × *Pre-stimulus pupil diameter*) + *β*_4_ · *Congruence* + *β*_5_ · *Object type* + *β*_6_ · (*Congruence* × *Object type*) + *Participant*. All regressors were z-scored.

|  | **Model 1** | **Model 2** |
| --- | --- | --- |
| **Dependent variable** | Recognition success | Recognition success |
| **Regressors** | Age,<br>Pre-stimulus baseline diameter,<br>Age × Baseline diameter. | Age,<br>Pre-stimulus baseline diameter,<br>Age × Baseline diameter. |
| **Factors** | - | Stimulus congruence,<br>Stimulus object type,<br>Congruence × Object type. |
| **Random effect** | Participant | Participant |
| $R^2_{\text{marginal}}$ | 0.002 | 0.026 |
| $R^2_{\text{conditional}}$ | 0.061 | 0.086 |
| **AIC** | 5824.910 | 5749.060 |
| **BIC** | 5856.850 | 5800.156 |
| **Overdispersion factor** | 0.976 | 0.975 |
| **Optimizer** | bobyqa | bobyqa |
| $\beta_0$<br>(95% CI, SE, p) | **0.4025**<br>([0.276, 0.529], 0.0647, <**0.001**) | **0.4106**<br>([0.281, 0.540], 0.0661, <**0.001**) |
| $\beta_1$ | 0.0729<br>([-0.056, 0.202], 0.0656, 0.267) | 0.0821<br>([-0.049, 0.213], 0.0670, 0.221) |
| $\beta_2$ | -0.0441<br>([-0.143, 0.055], 0.0504, 0.382) | -0.0447<br>([-0.145, 0.056], 0.0511, 0.382) |
| $\beta_3$ | -0.0275<br>([-0.139, 0.083], 0.0566, 0.627) | -0.0355<br>([-0.148, 0.077], 0.0573, 0.535) |
| $\beta_4$ | - | **0.1808**<br>([0.056, 0.306], 0.0637, **0.005**) |
| $\beta_5$ | - | **-0.5445**<br>([-0.670, -0.420], 0.0638, <**0.001**) |
| $\beta_6$ | - | -0.0594<br>([-0.309, 0.190], 0.1273, 0.641) |
| *Random effect*<br>($\sigma^2$, 95% CI, ICC) | (0.204, [0.128, 0.326], 0.059) | (0.216, [0.136, 0.344], 0.062) |

**Table S4:** Comparison of model parameters, model diagnostics, regression estimates and associated p-values, between two implemented generalized linear mixed models. **Model 1** is given by the following formula: logit(ℙ[*Recognition success* = 1]) = *β*_0_ + *β*_1_ · *Age* + *β*_2_ · *Maximum dilation* + *β*_3_ · (*Age* × *Maximum dilation*) + *Participant*. Formula of **model 2** is: logit(ℙ[*Recognition success* = 1]) = *β*_0_ + *β*_1_ · *Age* + *β*_2_ · *Maximum dilation* + *β*_3_ · (*Age* × *Maximum dilation*) + *β*_4_ · *Congruence* + *β*_5_ · *Object type* + *β*_6_ · (*Congruence* × *Object type*) + *Participant*. All regressors were z-scored.

|  | Model 1 | Model 2 |
| --- | --- | --- |
| **Dependent variable** | Recognition success | Recognition success |
| **Regressors** | Age,<br>Maximum dilation,<br>Age × Maximum dilation. | Age,<br>Maximum dilation,<br>Age × Maximum dilation. |
| **Factors** | - | Stimulus congruence,<br>Stimulus object type,<br>Congruence × Object type. |
| **Random effect** | Participant | Participant |
| $R^2_{\text{marginal}}$ | 0.003 | 0.025 |
| $R^2_{\text{conditional}}$ | 0.061 | 0.085 |
| **AIC** | 4804.640 | 4745.670 |
| **BIC** | 4835.623 | 4795.252 |
| **Overdispersion factor** | 0.973 | 0.972 |
| **Optimizer** | bobyqa | bobyqa |
| $\beta_0$<br>(95% CI, SE, p) | **0.4161**<br>([0.282, 0.550], 0.0683, <**0.001**) | **0.4232**<br>([0.286, 0.560], 0.0699, <**0.001**) |
| $\beta_1$ | 0.0980<br>([-0.037, 0.233], 0.0690, 0.156) | 0.0985<br>([-0.040, 0.237], 0.0707, 0.163) |
| $\beta_2$ | 0.0215<br>([-0.060, 0.103], 0.0415, 0.605) | 0.0104<br>([-0.072, 0.093], 0.0421, 0.804) |
| $\beta_3$ | -0.0349<br>([-0.126, 0.056], 0.0466, 0.454) | -0.0262<br>([-0.119, 0.066], 0.0471, 0.578) |
| $\beta_4$ | - | **0.1700**<br>([0.032, 0.308], 0.0702, **0.015**) |
| $\beta_5$ | - | **-0.5397**<br>([-0.678, -0.402], 0.0704, <**0.001**) |
| $\beta_6$ | - | -0.0306<br>([-0.306, 0.245], 0.1407, 0.828) |
| *Random effect*<br>($\sigma^2$, 95% CI, ICC) | (0.202, [0.122, 0.333], 0.058) | (0.215, [0.130, 0.351], 0.061) |

**Table S5:** Comparison of model parameters, model diagnostics, regression estimates and associated p-values, between two implemented linear mixed models. **Model 1** is given by the following formula: *Dilation amplitude* = *β*_0_ + *β*_1_ · *Congruence* + *β*_2_ · *Object type* + *β*_3_ · (*Congruence* × *Object type*) + *Participant*. Formula of **model 2** is: *Dilation amplitude* = *β*_0_ +*β*_1_ ·*Congruence*+*β*_2_ ·*Object type*+*β*_3_ ·(*Congruence*×*Object type*)+*Participant*+*Stimuli*. Post-hoc tests are reported with corrected p-values. The dependent variable was z-scored.

|  | Model 1 | Model 2 |
| --- | --- | --- |
| **Dependent variable** | Dilation amplitude | Dilation amplitude |
| **Regressors** | - | - |
| **Factors** | Stimulus congruence,<br>Stimulus object type,<br>Congruence × Object type. | Stimulus congruence,<br>Stimulus object type,<br>Congruence × Object type. |
| **Random effects** | Participant | Participant,<br>Stimulus. |
| $R^2_{marginal}$ | 0.005 | 0.004 |
| $R^2_{conditional}$ | 0.283 | 0.289 |
| **AIC** | 14314.201 | 14309.542 |
| **BIC** | 14373.886 | 14374.341 |
| **Overdispersion factor** | N/A | N/A |
| **Optimizer** | bobyqa | bobyqa |
| $\beta_0$ | 0.0314 | 0.0316 |
| (95% CI, SE, p) | ([-0.081, 0.144], 0.057, 0.585) | ([-0.082, 0.145], 0.058, 0.587) |
| $\beta_1$ | 0.0397 | 0.0341 |
|  | ([-0.006, 0.085], 0.023, 0.086) | ([-0.021, 0.089], 0.028, 0.224) |
| $\beta_2$ | **-0.0697** | **-0.0658** |
|  | ([-0.115, -0.024], 0.023, **0.003**) | ([-0.121, -0.011], 0.028, **0.020**) |
| $\beta_3$ | **-0.2199** | **-0.2177** |
|  | ([-0.311, -0.129], 0.046, **<0.001**) | ([-0.327, 0.108], 0.056, **<0.001**) |
| *Random effects*<br>($\sigma^2$, 95% CI, ICC) | (0.287, [0.211, 0.390], 0.279) | (0.287, [0.212, 0.390], 0.281)<br>(0.007, [0.001, 0.014], 0.009) |
| **Post-hoc tests** |  |  |
| Comparison (t, p) |  |  |
| Congruent - Animal vs.<br>Congruent - Furniture | (-1.222, 0.266) | (-1.087, 0.336) |
| Congruent - Animal vs.<br>Incongruent - Animal | **(-4.575, <0.001)** | **(-3.622, <0.001)** |
| Congruent - Animal vs.<br>Incongruent - Furniture | (0.915, 0.360) | (0.802, 0.424) |
| Congruent - Furniture vs.<br>Incongruent - Furniture | (2.138, 0.050) | (1.891, 0.092) |
| Incongruent - Animal vs.<br>Congruent - Furniture | **(3.336, <0.001)** | **(2.529, 0.026)** |
| Incongruent - Animal vs.<br>Incongruent - Furniture | **(5.504, <0.001)** | **(4.434, <0.001)** |

